# Gradual change of cortical representations with growing visual expertise for synthetic shapes

**DOI:** 10.1101/2023.01.24.525322

**Authors:** Ehsan Kakaei, Jochen Braun

## Abstract

Visual expertise for particular categories of objects (*e*.*g*., birds, mushrooms, minerals, and so on) is known to enhance cortical responses in parts of the ventral occipitotemporal cortex. How is such additional expertise integrated into the prior cortical representation of life-long cumulative visual experience? To address this question, we record multivariate BOLD responses to synthetic visual objects and track changes in pairwise distance as initially unfamiliar objects gradually become familiar (‘representational similarity analysis’, RSA).

We find view-independent responses to synthetic shapes (*i*.*e*., an invariant representation) in large parts of the ventral occipitotemporal cortex, including the primary visual cortex. Surprisingly, the quality of representation is already high even when shapes are still unfamiliar. As shapes become familiar with repeated viewing, this quality remains high, but representational geometry shifts with more experience. For visually similar shapes that are not repeated and therefore remain unfamiliar, the representation degrades and becomes progressively marginalized.

We conclude that, at least for highly dissimilar synthetic shapes, functional imaging responses can reveal detailed representational geometry at the level of exemplars, as well as gradual changes in this geometry due to experience and learning.

## 1 Introduction

An essential aspect of visual object recognition is the processing of visual shape. The neural substrate of shape processing includes the ventral visual pathway, which in humans extends over the ventral occipitotemporal cortex from the occipital pole to the lateral occipital cortex, fusiform gyrus, and beyond (reviewed by Grill-Spector and Weiner 2014; Bi et al. 2016; Kravitz et al. 2013; Weiner and Zilles 2016). Functional imaging studies of ventral occipitotemporal cortex reveal intriguing functional anatomy, with responsiveness to specific object categories (*e*.*g*., faces, scenes, body parts) changing systematically over the cortical surface along several large-scale anatomical gradients (*e*.*g*., animate-inanimate, large-small, feature-whole, or perception-action; Grill-Spector et al. 2004; Konkle and Oliva 2012; Grill-Spector and Weiner 2014; Freud et al. 2017; Yildirim et al. 2019; Wurm and Caramazza 2021).

Experience and learning improve object recognition performance, and also modify shape processing in the ventral occipitotemporal cortex. Indeed, functional imaging evidence shows that particular visual expertise – being able to identify and categorize visually similar objects of a particular kind – often entails moderate but anatomically distributed changes in the pre-existing responsiveness to shape (reviewed by Bukach et al. 2006; de Beeck and Baker 2010; Harel et al. 2013; Gauthier and Tarr 2016). This has been established by comparing novices and experts for identifying particular categories of natural objects (*e*.*g*., birds, mushrooms, minerals, degraded images; McGugin et al. 2012; Roth and Zohary 2015; Connolly et al. 2012; Freud et al. 2017; Martens et al. 2018; Cetron et al. 2019; Duyck et al. 2021), as well as by comparing observers before and after they have learned to categorize initially unfamiliar synthetic shapes (*e*.*g*., computer-generated ‘greebles’, ‘spikies’, or ‘ziggerins’; Gauthier et al. 1999; de Beeck et al. 2006; Yue et al. 2006; Wong et al. 2009; Brants et al. 2011; Wong et al. 2012).

Here we map the cortical representation of synthetic visual objects and track gradual changes as initially unfamiliar objects become progressively familiar with learning. We wondered how pre-existing shape representations would accommodate and integrate novel synthetic objects. We further wondered whether representational changes would be specific to learned objects or extend also to other objects of the same kind. Finally, we wondered how representational changes would be distributed anatomically in the ventral occipitotemporal cortex. To address these questions, we analyzed ‘representational similarity’ of spatiotemporal BOLD patterns (Kriegeskorte et al., 2008 b; Haxby, 2012), which offers a potentially sensitive measure for the information encoded in neural activity and may also be related to similarity as perceived by human observers (Charest and Kriegeskorte, 2015; Nestor et al., 2016; Collins and Behrmann, 2020).

Most previous studies of visual expertise identified cortical sites associated with a particular object category by comparing BOLD activity either between novices and experts or before and after learning. We extend this work in three ways: firstly, by analyzing the representational geometry revealed by multivariate BOLD activity, secondly, by establishing representational distance at the level of object exemplars rather than object categories and, thirdly, by monitoring gradual changes over time, as observers gain familiarity with a particular set of object exemplars. Few previous studies have attempted to resolve shape representations in comparable detail (Eger et al., 2008; Brants et al., 2016; Visconti di Oleggio Castello et al., 2021; Duyck et al., 2021). To progress fine-grained analysis of representational geometry, we developed synthetic shapes for which visual expertise is acquired comparatively slowly (Kakaei et al., 2021) and took advantage of a numerically tractable method for linear discriminant analysis in O(1000) dimensions (DLDA; Yu and Yang 2001).

Our results show view-invariant representations of shape over surprisingly extensive regions of the ventral occipitotemporal cortex, including the fusiform gyrus, lateral occipital areas, and primary visual cortex. Representational distance between recurring exemplars is initially high and remains largely stable during learning, suggesting that newly familiar shapes are accommodated by and encoded within pre-existing representations. In contrast, non-recurring exemplars (which are necessarily unfamiliar) substantially decrease distances between each other and increase distances to recurring exemplars, implying a progressive marginalization in representational space. The distinction between recurring and non-recurring objects, which is material to the behavioral task of observers, also becomes pronounced in certain areas of the frontal and parietal cortex.

## 2 Methods

### 2.1 Participants

Eight healthy participants (4 female and 4 male; age 25 to 32 years) with normal or corrected to normal vision, took part in the functional imaging experiment. Twelve additional participants took part in purely behavioral experiments of almost identical design reported previously (Kakaei et al., 2021). All participants were paid and gave informed consent. Ethical approval was granted under Chiffre 30/21 by the ethics committee of the Faculty of Medicine of the Otto-von-Guericke University, Magdeburg.

### 2.2 Experimental Paradigm

Our goal was to investigate the learning of visual object recognition. To this end, complex three-dimensional objects were computer-generated and presented as described previously (Kakaei et al., 2021). Briefly, all objects were highly characteristic and dissimilar from each other. Fifteen objects recurred at least 190 (mean±S.D: 216 ± 9) times each over the course of the experiment (‘recurring’ objects), whereas 360 other objects appeared exactly once (‘non-recurring’ or ‘singular’ objects) (**Fig. 1A**). Each object was presented for 3*s* from a random point of view while rotating slowly clock-wise or counterclock-wise(144^°^/*s* or 0.4*Hz*) around a randomly oriented axis with an inclination of 0^°^, 45^°^ or −45^°^, in the frontal plane (**Fig. 1B**). During the object presentation, observers responded to each object by classifying it as either ‘familiar’ or ‘unfamiliar’ by means of a button press. (The responding hand was counter-balanced over observers and conditions). Over the course of the experiment, all observers gradually became familiar with the ‘recurring objects’ (see below). The average time-course of learning, as established by a simplified signal detection and reaction-time (RT) analysis, is illustrated in **Fig. 1C**. All stimuli were generated with MATLAB (The MathWorks, Inc.) and presented with the psychophysics toolbox (Brainard, 1997).

**Figure 1:**
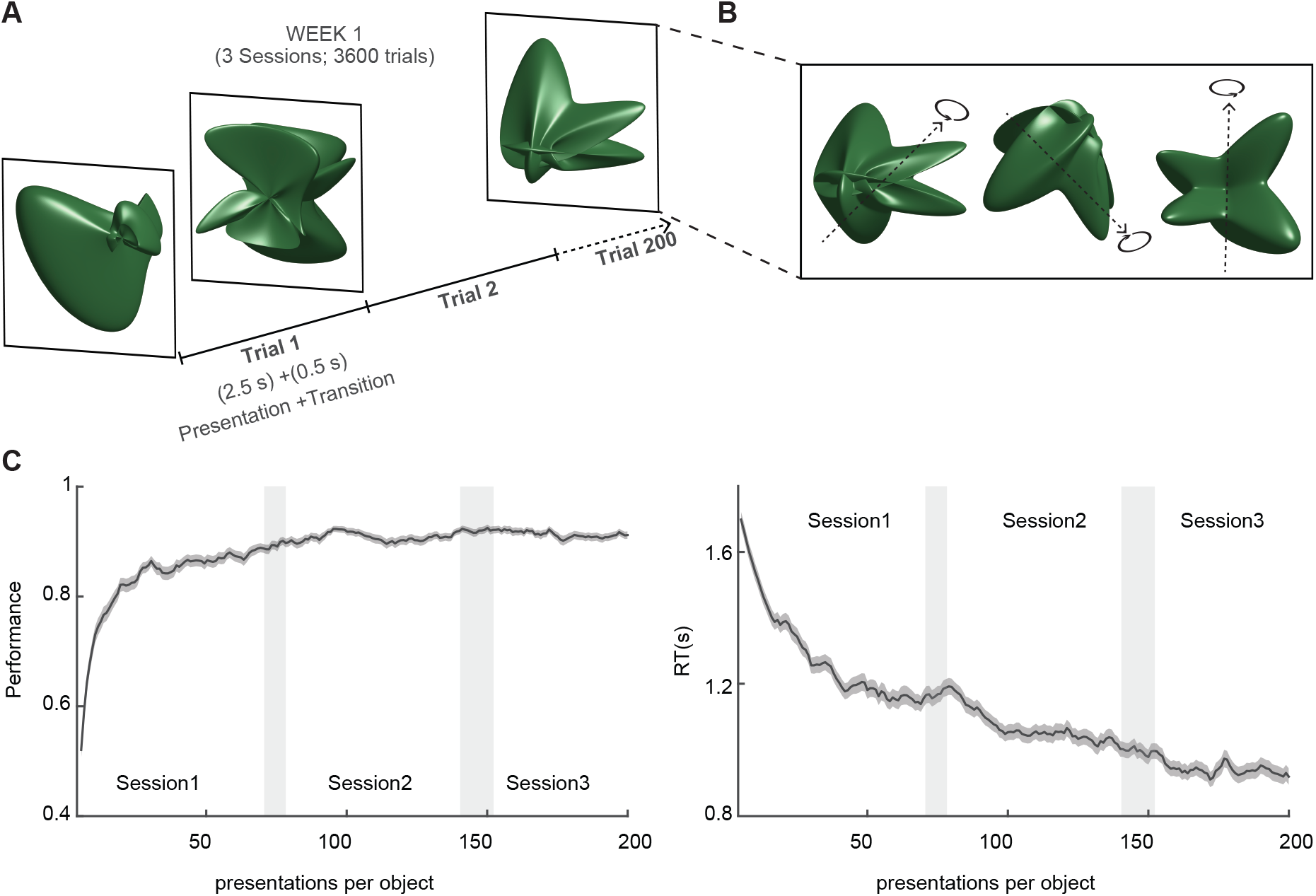
Experimental paradigm of the experiment. **A)** Complex 3D objects, rotating slowly, were shown in sequences with 200 object presentations (180 recurring and 20 non-recurring presentations) of 3 seconds long (2.5s presentation + 0.5s transition). Each week, observers participated in 18 such runs over 3 consecutive sessions. During each trial, observers were asked to categorize each object as either ‘familiar’ or ‘unfamiliar’ by button presses. **B)** Objects appeared with a random initial phase and rotated clockwise or counter-clockwise around an axis with an inclination of 0^°^, 45^°^, or *−*45^°^. **C)**Over the course of the experiment, as subjects learned to distinguish recurring objects from non-recurring ones, performance increased and reaction times (RT) decreased. Adapted from Kakaei et al. 2021.

Presentation sequences started with a random recurring object and immediate repetitions (*X* → *X*) and direct returns (*X* → *Y* → *X*) were not allowed. All sequences comprised exactly 200 objects and were selected such as to ensure that every recurring object appeared comparably often over the course of the experiment. Objects were presented either in ‘unstructured’ sequences without sequential dependencies or in ‘strongly structured’ sequences with sequential dependencies (Kakaei et al., 2021). However, this difference is irrelevant in the present context and the effects of sequence structure will be described elsewhere. Of 200 objects, 180 objects were recurring, with each being visited 12 ±1.9 times during a given sequence. The remaining 20 objects were non-recurring and were interspersed at random intervals. The total duration of each sequence was 600 *s*.

In the first week of the experiment, participants experienced one set of fifteen recurring objects in one type of sequence whereas, in the third week, they experienced another set of fifteen objects in the other type of sequence. In each week, participants performed in three fMRI sessions (on three separate days), each time viewing six sequences with 200 object. The second week of the experiment was free.

Prior to the fMRI sessions, participants familiarized themselves with the task design in a ‘sham experiment’ of a similar design. Specifically, they viewed 6 sequences of two-dimensional objects (recurring and non-recurring) during a single session and tried to classify these objects as ‘familiar’ or ‘unfamiliar’. After each week with fMRI sessions, participants performed a spatial search task with recurring and non-recurring objects to confirm that every participant had in fact become familiar with each of the recurring objects that they had viewed some 200 times during the preceding week (Kakaei et al., 2021).

### 2.3 MRI acquisition

All magnetic-resonance images were acquired on a 3T Siemens Prisma scanner with a 64-channel head coil. Structural images were T1-weighted sequences (**MPRAGE** TR = 2500 ms, TE = 2.82 ms, TI = 1100 ms, 7^°^ flip angle, isotropic resolution 1 × 1 × 1 *mm* and matrix size of 256 × 256 × 192). Functional images were T2*-weighted sequences (TR = 1000 ms, TE = 30ms, 65^°^ flip angle, resolution of 3 × 3 × 3.6 *mm* and matrix size of 72 × 72 × 36). Field maps were obtained by gradient dual-echo sequences (TR = 720 ms, TE1 = 4.92 ms, TE2 = 7.38 ms, resolution of 1.594 × 1.594 × 2 *mm* and matrix size of 138 × 138 × 72).

### 2.4 fMRI pre-processing

The fMRI pre-processing procedure was similar to that published previously (Dornas and Braun, 2018). First, DICOM files were converted into NIFTI format using MRICRON (MRICRON Toolbox, Maryland, USA, NIH). Then, brain tissues were extracted and segmented using BET (Smith, 2002) and FAST (Zhang et al., 2001). Field map correction, head motion correction, spatial smoothing, high-pass temporal filtering, and registration to structural and standard images were performed with the MELODIC package of FSL (Beckmann and Smith, 2004).

Field map correction and registration to structural image were carried out using Boundary-Based Registration (BBR; Greve and Fischl 2009). MELODIC uses MCFLIRT (Jenkinson et al., 2002) to correct for head motion. Spatial smoothing was performed with SUSAN (Smith and Brady, 1997), with full width at half maximum set at FWHM= 5 *mm*. To remove low-frequency artefacts, we applied a high-pass filter of the cut-off frequency *f* = 0.01Hz, *i*.*e*., oscillations/events with periods of more than 100 seconds were removed. To register the structural image to Montreal MNI152 standard space with isotropic 2 *mm* voxel size, we used FLIRT(FMRIB’s Linear Image Registration Tool; Jenkinson and Smith 2001; Jenkinson et al. 2002) with 12 degrees of freedom (DOF) and FNIRT (FMRIB’s Nonlinear Image Registration Tool) to apply the non-linear registration. To further reduce artifacts arising from head motion, we applied despiking with a threshold of *λ* = 100 using BrainWavelet toolbox (Patel et al., 2014). Later, we regressed out the mean CSF activity as well as 12 DOF translation and rotation factors predicted by a motion correction algorithm (MCFLIRT). Afterward, the time series of each voxel was whitened and detrended.

Finally, the 160, 099 voxels of MNI152 space were grouped into 758 functional parcels according to the MD758 atlas (Dornas and Braun, 2018). Each functional parcel is associated with an anatomically labeled region of the AAL atlas (Tzourio-Mazoyer et al., 2002) and comprises approximately 200 voxels or approximately 1.7*cm*^*3*^ of gray matter volume (212 ± 70 voxels, range 45 to 462 voxels). This parcellation offers superior cluster quality, correlational structure, sparseness, and consistency with fiber tracking, compared to other parcellations of similar resolution (Dornas and Braun, 2018). In the present context, its main advantage is the consistency of activity patterns within functional parcels. In a previous study of resting state activity, we found that voxel-by-voxel correlations within functional parcels were highly similar in different scanning sessions. To a lesser extent, this was the case even for different observers (Dornas and Braun, 2018). Accordingly, we hypothesized that voxel-by-voxel activity within functional parcels might provide consistent information also about task states.

### 2.5 fMRI data analysis

In every functional parcel, the activity pattern during and following an object presentation was recorded over 9 *s* (from 2 to 11 *s* after object onset). For a parcel with *N*_*vox*_ voxels, such an activity pattern may be represented as a 9 · *N*_*vox*_ -dimensional vector (**Fig. 2.A**). To investigate the separability of activity patterns associated with different objects, we employed a representational similarity analysis (Kriegeskorte et al., 2008 a) to identify parcels with significant selectivity for individual recurring objects (**Fig. 2B**). In object-identity-selective parcels, we examined the separability of object representations over the course of learning, as well as the separability of representations of recurring and non-recurring objects.

**Figure 2:**
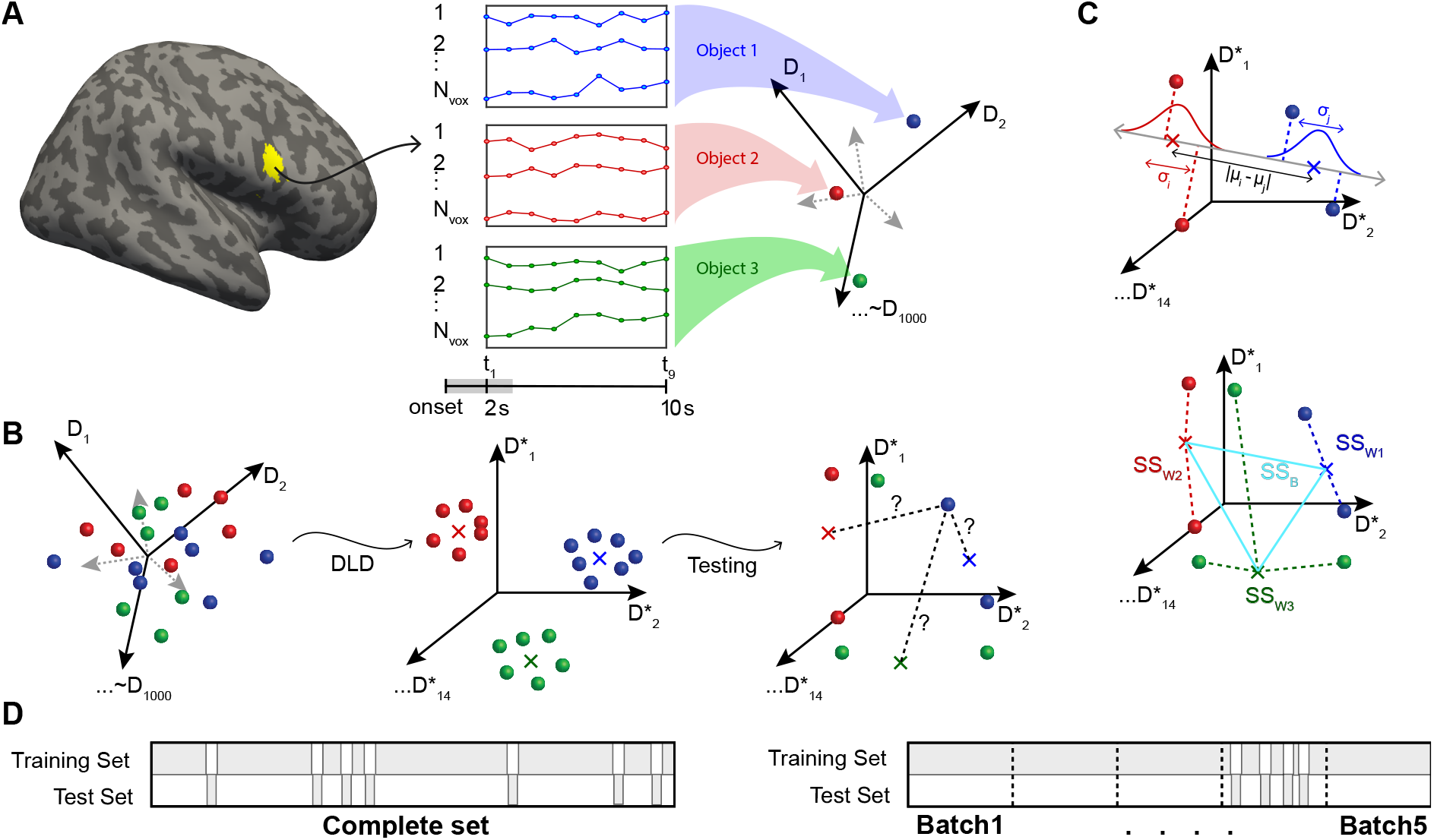
Analysis of fMRI activity with direct linear discriminant analysis, or DLDA. For each functional parcel, DLDA identified the 14-dimensional space that optimally discriminated the 15 classes of activity patterns associated with 15 *recurring* objects. Other activity patterns, such as those associated with *nonrecurring* objects, were also analyzed in this space. **A)** For a given parcel with *N*_*vox*_ voxels (*e*.*g*., yellow region Frontal-Inf-R-8), activity was recorded over 9 *s* during and following object presentation (2 to 11 *s* after onset). Each such activity pattern corresponds to a point (or vector) in a 9 *·N*_*vox*_ -dimensional space (right). Here, activity patterns associated with three object presentations are represented schematically (red, green, and blue spheres). **B)** To cross-validate discriminability, recurrent object presentations were divided randomly into a training set (90%) and a test set (10%). From the training set, the DLD subspace S was established. Here, exemplars (solid spheres) and class centroids (crosses) are represented schematically. Next, the projections into this space of test set patterns were compared to class centroids. **C)** Projection onto the line connecting class centroids *i* and *j* revealed the pairwise discriminability/dissimilarity *δ*_*i,j*_ of object classes *i* and *j* (top), and the distances to class centroids yielded the within-class and between-class variance of representations, *SS*_*W*_ and *SS*_*B*_, and the associated variance ratio *F* = *SS*_*B*_ */SS*_*W*_ (bottom). Additionally, a matrix of (mis-)classification probabilities *P* (*reported i* |*true j*) (a.k.a confusion matrix) could be obtained (not shown). **D)** To assess object representation generally, test presentations were drawn randomly from the complete set of object presentations (left). To assess changes over the duration of the experiment, the set of presentations was divided into five successive ‘batches’ and test presentations were drawn from one of these batches (bottom). In either case, the training set comprised all remaining presentations (*i*.*e*., the complement of the test set).

When analyzing the representation of object ‘identity’ and object ‘novelty’, we combined data from strongly structured and unstructured sequences, raising the effective number of observers to 16.

#### 2.5.1 Linear discriminant analysis

Representational similarity/dissimilarity is commonly analyzed in terms of the Euclidean, standardized Euclidean, or Mahalanobis distance between observed activity patterns in a high-dimensional space (Kriegeskorte and Diedrichsen, 2019). In the present case, the dimensionality of activity patterns in a functional parcel was the number of voxels *N*_*vox*_ in the parcel, times the number of time-points, *N*_*t*_ = 9 (9 *TR* of 1 *s*). Accordingly, the dimensionality was *N*_*dim*_ = *N*_*vox*_ × *N*_*t*_ = *O*(1000), with mean and standard deviation of *N*_*dim*_ = 1911±634 and range 405 to 4158. To analyze distance in this space, we applied Fisher’s Linear Discriminant Analysis (LDA) for multiple classes to perform a ‘supervised’ principal component analysis and to identify the (at most) (*κ* − 1)-dimensional subspace S that optimally discriminates *κ* = 15 classes of activity patterns. Optimality is defined here as simultaneously minimizing within-class variance and maximizing the between-class variance of activity patterns. A numerically tractable procedure for identifying this (*κ*−1)-dimensional subspace is available in terms of ‘direct LDA’ or DLDA (Yu and Yang, 2001; Ye et al., 2006). Briefly, this method first diagonalizes between-class variance to identify *κ* − 1 discriminative eigenvectors with non-zero eigenvalues, next diagonalizes within-class variance, and finally yields a rectangular matrix for projecting activity patterns from the original activity space (dimensionality *N*_*dim*_) to the maximally discriminative subspace S (dimensionality *κ* − 1 = 14) and back. As this method is linear and relies on all available degrees of freedom, its results are deterministic. Specifically, it results in dimensionally reduced activity patterns *x*_*jk*_, where *j* ∈ {1, …, 14} indexes (reduced) dimensions and *k* indexes trials. A Matlab implementation of this procedure is available on github.com/cognitive-biology/DLDA.

#### 2.5.2 Amplitudes, distances, and correlations

Activity patterns *x*_*jk*_ associated with trials *k* were analyzed in the maximally discriminative subspace 𝕊. The normalized amplitude 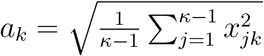 of such patterns exhibited an average value of ⟨*a*⟩ = 0.99. The normalized distance 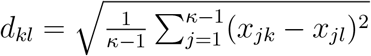 between patterns associated with trials *k* and *l* measured on average ⟨ *d* ⟩ = 1.40, consistent with distance expected between random patterns of this amplitude.

The pattern associated with successive trials exhibited a weak temporal correlation, with approximately 5% smaller distances at delays below 4 trials and approximately 2% larger distances at delays ranging from 6 to 15 trials (see **Figure S1** for details).

For certain analyses (Sections 2.5.7 and 2.5.10), we established for each parcel *w* the average delay-dependent distance *T*_*w*_(Δ*i*) = *d*_*w,u,r*_(Δ*i*) _*u,r*_ between patterns with relative delay Δ*i*, where the average was taken over subjects *u* and runs *r*. The timecourse *T*_*w*_ allowed us to discount temporal correlations by computing 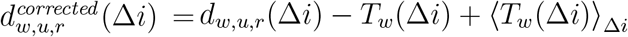, where ⟨ *T*_*w*_(Δ*i*) ⟩_Δ*i*_; is the average value over delays.

#### 2.5.3 Representation ‘identity’ for recurrent objects

Our observations comprised approximately 200 activity patterns for each of the 15 recurring object classes (for every observer and condition). To allow for cross-validation, we randomly divided these patterns in a larger ‘training set’ (90% or 190 ± 7.7 per object class) and a smaller test set (10% or 22 ± 0.9 per object class) (**Fig. 2B**). Note that the ‘training set’ comprised exclusively activity patterns associated with recurring objects. To reduce variance deriving from the random selection of test sets, this random selection was repeated *N*_*r*_ = 20 times and all statistical measures described below represent the average over all repetitions. As illustrated in **Fig. 2C**, in the discriminative subspace 𝕊, we compared the *n*_*i*_ pattern exemplars *x*_*ki*_ (where *k* = 1, …, *n*_*i*_) from the test set *I* to the class centroids 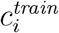 established for the training set *i* or, alternatively, the class centroids 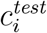 of the test set.

Several alternative approaches were used for this comparison. Firstly, the nearest class centroid 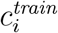 to each exemplar *x*_*k*_ was identified to establish a matrix of classification probabilities *P* (*j*|*i*) (probability that an exemplar of class *i* is nearest to the centroid of class *j*), also known as ‘confusion matrix’, as well as ‘classification accuracy’ *a* = ∑_*i*_ *P* (*i*|*i*)*P* (*i*), which is the probability that the nearest centroid is indeed of the correct object class.

Secondly, for each pair of object classes (*i, j*), object exemplars *x*_*ki*_ and *x*_*kj*_ from the test set were projected onto the line connecting the two test set centroids, 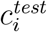 and 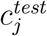, and a pairwise discriminability/dissimilarity / Mahalanobis distance *δ*_*i,j*_ was computed from the means, *μ*_*i*_ and *μ*_*j*_, and variances, 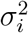 and 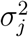, of these projections, as 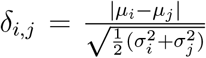. The average over all pairs of object classes was computed as 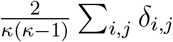.

Thirdly, given class centroids 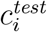 and overall centroid 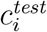, we computed the Euclidean distances 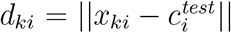 between exemplars *x*_*ki*_ and class centroid 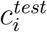 and, for each object class *i*, the ‘sum of squares’ as 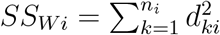. The ‘within-class’ variance of all classes was computed as 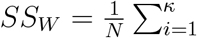, where 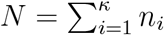. Similarly, from the Euclidean distances 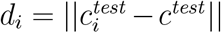 between individual and overall centroids, we computed ‘between-class’ variance 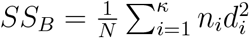. From the Euclidean distances 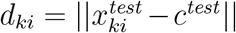 between exemplars and overall centroid, we computed ‘total’ variance 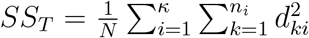. Variances *SS*_*W*_, *SS*_*B*_, and *SS*_*T*_ are also denoted, respectively, *SS*_*same*_, *SS*_*diff*_, and *SS*_*fam*_ further below. To quantify the discriminability of classes, a variance ratio was computed as *F*_*identity*_ = *SS*_*B*_(*N* − *κ*)*/SS*_*W*_ (*κ* − 1) (Anderson, 2001). The average within-class and between-class dispersion per dimension could be estimated as 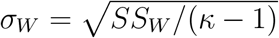 and 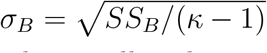, respectively.

This analysis was performed over all 8 observers and both conditions (types of presentation sequences), raising the effective number of observers to 16.

#### 2.5.4 Prevalence analysis

To test for a statistically significant representation of object identity in a functional parcel, we used ‘information prevalence inference’ as described by Allefeld et al. 2016. Briefly, we computed average classification performance (e.g., classification accuracy *a*_*obs*_ or f-ratio *F*_*obs*_) over *N*_*r*_ test sets, as well as over 10^3^ first-level permutations of object identities (in each of the *N*_*r*_ test sets). In principle, this could have been used to test an ‘individual null’ hypothesis, namely, the probability of obtaining the observed performance *a*_*obs*_ (or *F*_*obs*_) purely by chance. Instead, we computed the ‘minimum statistic’ *m*_*obs*_ = *min*_*k*_ *a*_*k*_ (or *m*_*obs*_ = *min*_*k*_ *F*_*k*_) over observers *k*, as well as over 10^5^ second-level permutations (drawn randomly from the first level permutations) and tested the ‘global null’ hypothesis, namely, the probability *p*_0_(*m*_*obs*_) of obtaining the observed minimum performance *m*_*obs*_ purely by chance. When this hypothesis could be rejected, we could infer statistically significant classification performance in at least *some* observers. To correct for multiple comparisons, we calculated the maximum over parcel for each of the second-level permutations and compared the observed minimum statistic *m*_*obs*_ against this distribution.

To establish the prevalence of classification performance over observers, we tested the ‘prevalence null’ hypothesis that the prevalence fraction *γ*_*true*_ of significant performance is smaller than, or equal to, a threshold *γ*. The largest value of *γ* for which this hypothesis can be rejected was then computed from 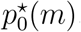, the number of observers *n* = 16, and the corrected significance threshold 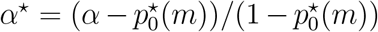 for a chosen threshold *α* = 0.05 (Allefeld et al., 2016):

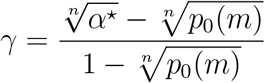

Any parcel with prevalence *γ* ≥ 0.5 (*i*.*e*., in which at least half of the observers exhibited an above-chance of effect) was considered to be ‘identity-selective’. A prevalence threshold *γ* ≥ 0.5 corresponds to a corrected probability 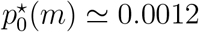 and a *minimal* accuracy near chance 6.67%.

#### 2.5.5 Representation of ‘novelty’ for non-recurring objects

Over the course of the experiment, observers learned to recognize recurring objects as ‘familiar’ and non-recurring objects as ‘novel’ **Fig. 1**. Accordingly, it is conceivable that neural representations of the class of 15 recurring objects are distinguishable from representations of the class of 360 non-recurring objects. To assess this possibility, we divided non-recurring and recurring objects into two sets of unequal size (approximately *N* = 216 × 15 recurrent or ‘familiar’ exemplars vs. *M* = 360 non-recurrent or ‘novel’ exemplars). From the Euclidean distances *d*_*k*_ = ||*x*_*k*_ − *c*|| between test set exemplars *x*_*k*_ and centroids 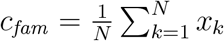 or 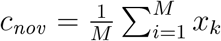, we obtained ‘within-class’ variance *SS*_*W*_ = *SS*_*fam*_ + *SS*_*nov*_, where 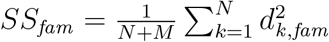 and 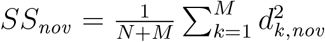. From distances *d*_*fam*_ = ||*c*_*fam*_ − *c*_*tot*_ || and *d*_*nov*_ = ||*c*_*nov*_ − *c*_*tot*_ || between class centroids and overall centroid 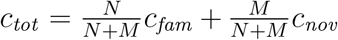, we obtained ‘between-class’ variance 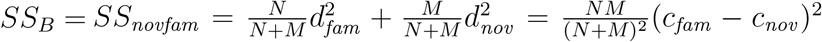. Finally, from distances *d*_*k*_ = ||*x*_*k*_ − *c*_*tot*_|| between exemplars and overall centroid, we obtained total variance 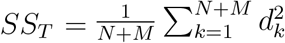. To quantify the discriminability of non-recurring and recurring objects, we formed the variance ratio *F*_*novelty*_ = *SS*_*B*_(*N* + *M* − 2)*/SS*_*W*_. Average within-class and between-class dispersion per dimension was obtained from 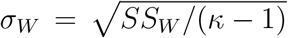 and 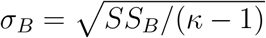, respectively.

#### 2.5.6 Changes with learning analyzed in ‘batches’

To assess the extent to which neural representations change over the course of learning, we divided all recurring object presentations into five successive ‘batches’ *B*_1_, *B*_2_, …, each with 20% of the presentations (**Fig. 2D**). In this way, we could select ‘test sets’ for crossvalidated DLDA from one particular batch, while retaining all other presentations as a ‘training set’. As every recurrent object was presented 210 ± 9 times over all sessions, a batch would comprise 42±1.8 presentations, a test set 21±0.9 and a training set 189±8.1 presentations. To reduce the variance deriving from test set selection, we repeated the random selection *N*_*r*_ = 20 times and averaged over repetitions.

To quantify representational changes over the course of learning, we computed the variance ratios 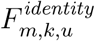 for each temporal window or batch *m*, identity-selective parcel *k*, and observer *u* ∈ {1, …, 16}. To summarize the changes in a particular identity-selective parcel *k*, we determined a ‘slope’ parameter 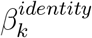 by fitting a linear mixed-model 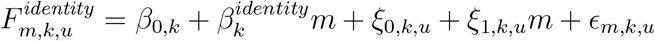 with subjects as grouping variables, where *β*_0,*k*_ was a fixed-effect coefficient, *ξ*_0,*k,u*_ and *ξ*_1,*k,u*_ were random effect coefficients, and ϵ was residual error.

To assess the statistical significance of the average ratio over observers, 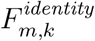, we shuffled (10^3^ permutations) the identity of recurring objects and established the distribution of variance ratios due to chance or data structure. With the mean *μ*_*m,k*_ and variance 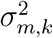of this distribution, we converted 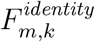 into z-score values 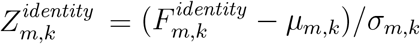.

Similarly, to assess the extent to which neural representations of novelty change over the course of learning, we divided all object presentations (recurring and non-recurring) into five successive ‘batches’ *B*_1_, *B*_2_, …, each with 20% of the presentations (**Fig. 2D**). Changes in the representation of object ‘novelty’ were assessed by fitting the ‘slope’ parameter 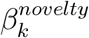 in a linear mixed-model 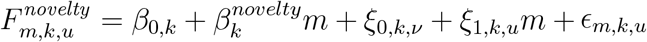, with subjects as grouping variables, where *β*_0,*i*_ was a fixed-effect coefficient, *ξ*_0,*k,u*_ and *ξ*_1,*k,u*_ were random effect coefficients, and ϵ was a residual error. To assess the significance of observer average ratios 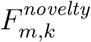, we calculated z-score values 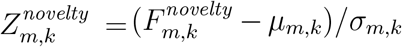 from the mean *μ*_*m,k*_ and variance 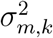 of the distribution obtained by shuffling object classification as ‘recurring’ or ‘non-recurring’ (10^3^ permutations).

An average learning trend *F*_*m*_ = ⟨*F*_*m,k,u*⟩ *k,u*_ was obtained (for both identity and novelty) in terms of slope parameter *β*_1_, by fitting a linear mixed-model *F*_*m,k,u*_ = *β*_0_ + *β*_1_*m* + *ξ*_0,*k,u*_ + *ξ*_1,*k,u*_*m* + ϵ _*m,k,u*_ with both parcels and subjects as grouping variables.

These analyses were performed over all 8 observers and both conditions (types of presentation sequences), raising the effective number of observers to 16.

#### 2.5.7 Geometry of identity and novelty representations

For the cross-validated analyses described above, activity patterns were divided randomly into training and test sets. A drawback of this approach was that the subspace 𝕊 changed with each training set. In order to be able to assess subtle differences in object representations during and between sessions, we repeated some analyses in a fixed subspace obtained from the complete set of all activity patterns *x*_*k*_. In this fixed subspace, we calculated the normalized amplitude 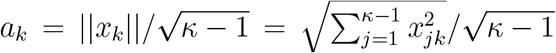 of individual patterns *k* and the normalized pairwise distance 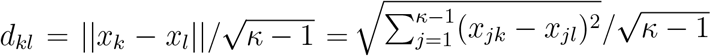 between two patterns *k* and *l*.

For each parcel *w*, observer *u*, and run *r*, we obtained the average amplitude 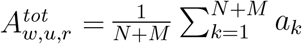 of all patterns, the average amplitude 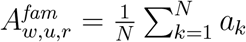 of *non-recurring* patterns, and the average amplitude 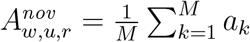 of *recurring* patterns. Similarly, we obtained the average pairwise distance 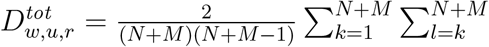 between all patterns, the average distance 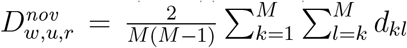 between non-recurring patterns, the average distance 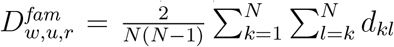 between recurring patterns, and the average distance 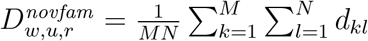 between pairs comprising one recurring and one non-recurring pattern. For recurring patterns, we further obtained the average distance 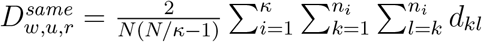 between pairs of recurring patterns in the *same* class and the average distance 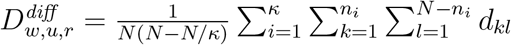 between pairs in *different* classes. All distances were corrected for the temporal autocorrelation by subtracting the time course of *T*_*w*_(*i, j*), as described above.

As described further above, the distances between individual activity patterns and different centroids – such as *c*_*tot*_, *c*_*nov*_, and *c*_*fam*_ – yielded total variance *SS*_*T*_ = *SS*_*tot*_, within-class variance *SS*_*W*_ = *SS*_*fam*_ +*SS*_*nov*_, and between-class variance *SS*_*B*_ = *SS*_*novfam*_. For recurring patterns, distances to individual class centroids *c*_*i*_ and overall centroid *c*_*fam*_ yielded total variance *SS*_*T*_ = *SS*_*fam*_, within-class variance *SS*_*W*_ = *SS*_*same*_ and between-class variance *SS*_*B*_ = *SS*_*diff*_.

These values were computed for each parcel *w*, observer *u*, and run *r*, in order to obtain variance fractions 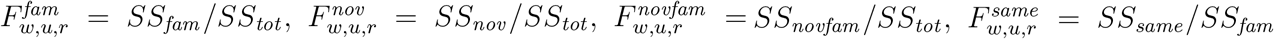, and 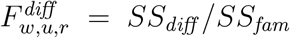, as well as variance ratios 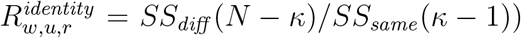 and 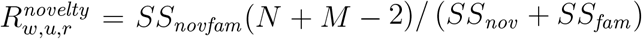.

#### 2.5.8 Stability of identity and novelty representations

To assess the stability of representations over the course of the experiment, we compared the average representation in individual runs *r* (centroids *C*_*r*_ of responses to exemplars) to the average representation over all runs (centroids *C*_*all*_). For subject *u*, identity-selective parcel *w*, and object class *i*, we calculated the Euclidean distance *D*_*u,k,i,r*_ between the relevant *C*_*r*_ and *C*_*all*_, and also the distance Δ*D*_*u,k,i,r*_ between the relevant centroids from successive runs, *C*_*r*_ and *C*_*r*+1_. After averaging over subjects *u*, identity-selective parcels *w*, and object classes *i*, we obtained *D*_*same*_ (*r*) and Δ*D*_*same*_ for recurring objects and by *D*_*nov*_ (*r*) and Δ*D*_*nov*_ (*r*) for non-recurring objects.

#### 2.5.9 Dimensional reduction

To visualize representational geometry, we randomly sampled 50 response patterns to each of the recurring and non-recurring objects within the first and the last sessions and calculated a 1600 × 1600 pair-wise distance matrix (*D*_*k,u*_) for each identity-selective parcel *k* and subject *u*. As we did not expect the activity patterns of different subjects to be comparable, we did not wish to directly average distance matrices over subjects. To circumvent this difficulty, we permuted the order of recurring objects 100 times and for each subject obtained an average matrix 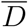 over permutations, which was then averaged over subjects. To visualize the representational geometry of identity in the first and the last session, we used multidimensional scaling (Matlab function *mdscale*, metric stress) to map the distances matrices for recurring objects (50 exemplars from the first session and 50 exemplars from the last session) into a two-dimensional space. To visualize the representational geometry of novelty, we restricted the distance matrix to non-recurring objects (50 exemplars from the first session and 50 exemplars from the last session) and just 3 of the 15 recurring objects (20 exemplars from either session).

#### 2.5.10 Changes with learning analysed by ‘runs’

To assess representational changes over the course of learning at greater temporal resolution, we computed average amplitudes *A*_*w,u,r*_, distances *D*_*w,u,r*_, variances *SS*_*w,u,r*_, and variance ratios *F*_*w,u,r*_, as described above, for each parcel *w*, observer *u* ∈ {1, …, 16}, and run *r*. Within each session *s*, we assessed the changes of these parameters *Y* ∈ {*A, D, SS, F* } over runs *r′* ∈ *s* by determining an ‘inclination’ parameter *β*_*s*_ for identityselective *k* and non-selective parcels *k*^*′*^. Each *β*_*s*_ coefficient was acquired from a linear mixed-model 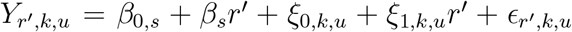 with subjects and parcels as grouping variables, where *β*_0,*s*_ was a fixed-effect coefficient, *ξ*_0,*k,u*_ and *ξ*_1,*k,u*_ were random effect coefficients, and ϵ was residual error. The same approach was used to assess gradual changes over runs in the centroid-to-centroid distances *D*_*same*_ (*r*), Δ*D*_*same*_ (*r*), *D*_*nov*_ (*r*), and Δ*D*_*nov*_ (*r*).

## 3 Results

### 3.1 Representation of object identity

As described in the Methods, we trained a DLD classifier to categorize objects in terms of identity and we used various measures – classification accuracy *a*_*w,u*_, average pairwise dissimilarity *δ*_*w,u*_, and between to within-class variance ratio *F*_*w,u*_ – to quantify the availability of ‘identity’ information in every parcel *w* and observer *u* ∈ {1, …, 16} (8 participants and 2 conditions). All three measures proved to be highly correlated and supported similar conclusions. A representative example is the correlation between classification accuracy *a*_*w,u*_ and variance ratio *F*_*w,u*_ (*ρ* = 0.94, *p* < 0.001), which is illustrated in **Fig. 3B**. The correlations between *a*_*w,u*_ and *δ*_*w,u*_ (*ρ* = 0.95, *p* < 0.001), and between *δ*_*w,u*_ and *F*_*w,u*_ (*ρ* = 0.98, *p* < 0.001) were comparably high. The accuracy *a*_*w,u*_ for both conditions (structured and unstructured sequences) was highly correlated as well, demonstrating test-retest consistency (**Fig. S2**).

**Figure 3:**
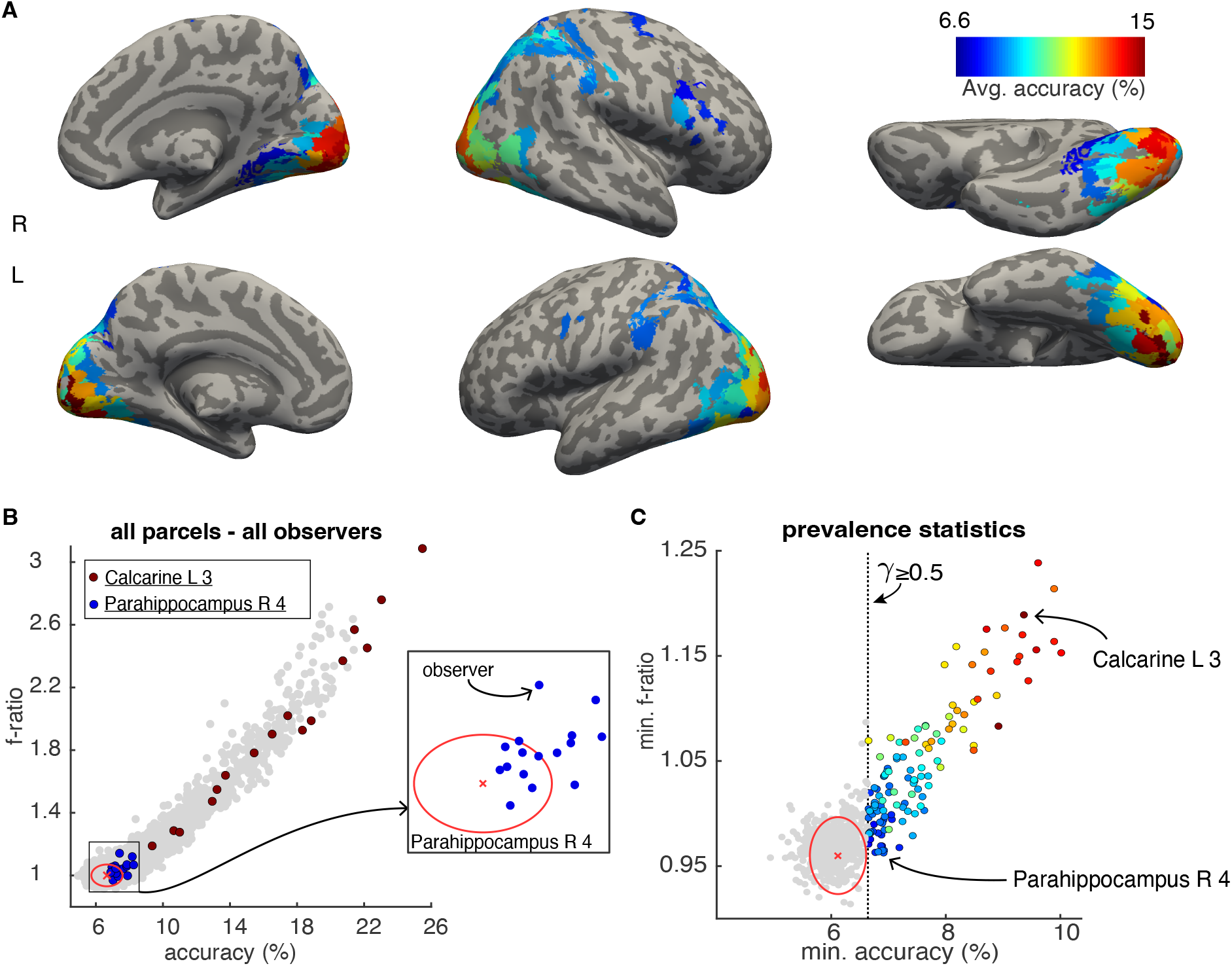
Neural representation of object identity. **A)** Classification accuracy based on neural responses of identity-selective parcels, average over observers *a*_*ν*_ The color scale ranges from chance to the maximal observed accuracy (6.67% to 17%). Altogether, 124 parcels proved identity-selective (prevalence *γ* ≥ 0.5), including 49 occipital, 17 fusiform/temporal, 22 parietal and 4 frontal parcels (Table 1). **B)** Classification accuracy *a*_*k,ν*_ and variance ratio *F*_*k,ν*_ for all observers *k* and parcels *ν*. Two particular parcels are highlighted (Calcarine-L-3 in red, Parahippocampus-R-4 in blue), to illustrate variability between individual observers. The distribution obtained with shuffled object identities are indicated as well (red cross and ellipse, representing mean ±3 S.D.). The Parahippocampus-R-4 values and the shuffle distribution are shown also magnified (inset). **C)** Prevalence statistics based on minimal values, taken over all observers. Minimal values 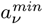 and 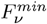 for all parcels. Identity-selective parcels are colored as in **A)**. The prevalence threshold *γ* ≥ 0.5 corresponds to a minimal accuracy near chance 6.67% (dotted vertical line). The distributions obtained with shuffled identities are indicated as well (red cross and ellipse).

Not unexpectedly, the variability between individual observers was considerable. Whereas a few parcels exhibited significant accuracy *a*_*w,u*_ and variance ratio *F*_*w,u*_ in all observers (e.g., Calcarine-L-3), in many parcels the representation of object identity was significant only in some observers (e.g., Parahippocampus-R-4).(**Fig. 3B**). To address this variability, we sought to identify parcels in which ‘identity’ was represented significantly in a majority of observers. To compute the ‘prevalence’ *γ* of significant representation among observers, we compared the minimal values over observers with the minimal values obtained from shuffled data (see Methods for details, red ellipse in **Fig. 3C**).

Out of 758 parcels, 124 were ‘identity-selective’ in that they exhibited a prevalence *γ* ≥ 0.5. Among these were 49 parcels in the occipital cortex, 17 in the fusiform or temporal cortex, 22 in the parietal cortincluding 49 occipitalex, and 4 in the frontal cortex. In general, we observed a pronounced posterior-anterior gradient. Whereas many parcels at the posterior pole of the brain exhibited high classification accuracy, this tendency diminished towards progressively more anterior locations (**Fig. 3.A, Table A.1, Fig. S3**). To better compare average accuracy between identity-selective parcels, we assigned such parcels to the 25 visual cortical regions defined by Wang et al. 2015, as well as to anterior inferior temporal cortex (AIT) and inferior frontal cortex (IFC) as two further regions. This summary of our results shows that accuracy was comparable in early visual areas (V1-hV4) and in the posterior-ventrolateral regions of the temporal lobe, whereas accuracy was lower in the anterior temporal cortex, the inferior frontal cortex, and in parietal cortical areas (**Fig. 8**).

Somewhat contrary to our expectations, identity-related representations were evident even in early visual cortical areas, which are thought to represent basic visual features such as orientation or motion, not just in higher areas of the ventral stream, which are thought to represent categorical object information (Grill-Spector and Weiner, 2014; Kravitz et al., 2013).

Apparently, the spatiotemporal BOLD activity elicited by visual objects encoded identity-specific information even though rotating objects were presented from varying points of view (i.e., random phase and axis of rotation). As identity encoding seemed impervious to this randomized presentation, we speculate that spatiotemporal activity may have reflected invariant relationships between multiple object features, rather than individual features that happened to be particularly salient.

Additionally, we observed identity-related representations also in ‘dorsal stream’ cortical areas, which are thought to encode goal-and task-related object features (Perry and Fallah, 2014; Christophel et al., 2017). Note, however, that any linear correlation between the respective spatiotemporal activations of cortical areas will result in some degree of sharing of identity information. Thus, whenever two areas are sufficiently interconnected to express spatiotemporally correlated activity patterns, they will share identity information to some degree.

### 3.2 Representational changes with learning

In order to assess changes in the neural representation of object identity over the course of learning, we established the ratio of between- and within-class variance in the maximally discriminate subspace S, for both object identity (15 classes formed by parcel responses to 15 recurring objects) and object novelty (2 classes formed by parcel responses to recurring and non-recurring objects, respectively) over five successive and non-overlapping periods of the experiment (termed ‘batches’, see Methods for details). These two variance ratios served as an index for the quality of the neural representation of ‘identity’ and ‘novelty’.

As direct linear discriminant analysis (DLDA) ‘whitens’ within-class variance to unity in each of the *κ* − 1 dimensions of optimally discriminant space S, the numerical value of ‘within-class’ variance *SS*_*W*_ was expected to be *κ* − 1 = 14. Indeed, the observed values for object ‘identity’, object ‘novelty’, and shuffled object identity/novelty were 14 ± 0.01. The numerical value of the ‘between-class’ variance *SS*_*B*_ was expected to be comparable or larger. Indeed, the values observed for object ‘identity’ was 18 ± 0.24 and for object ‘novelty’ was 27 ± 0.44. To obtain the average dispersion per dimension, within-class or between-class, these values were normalized with 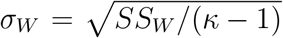 and 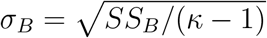, respectively.

The results were expressed in terms of z-score values 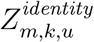 and 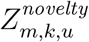 for identity-selective parcels *k*, and observers *u* ∈ {1, …, 16} and for successive batches *m* ∈ {1, …, 5} (8 participants and 2 conditions). The average over observers and identity-selective parcels, 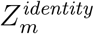 and 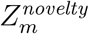, was highly significant for all batches (*p* < 0.001) (**Fig. 4**). Over successive batches, the representation of identity, as measured by 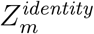, weakened slightly but significantly (*p* < 0.05). In contrast, the representation of novelty, as measured by 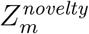, strengthened considerably, especially between batches *m* = 1 and *m* = 2 (*p* < 0.001).

**Figure 4:**
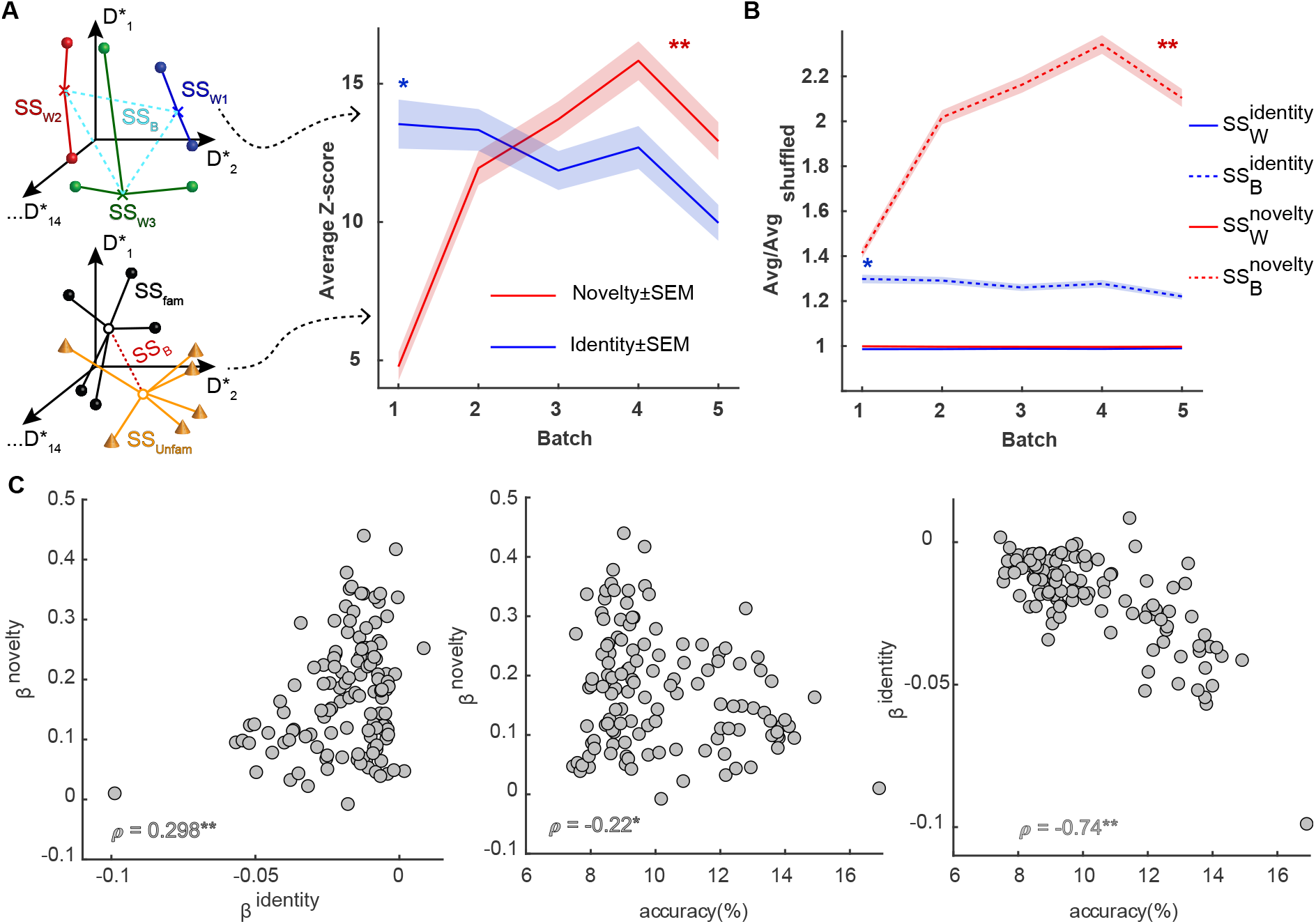
The representation of ‘identity’ and ‘novelty’ changes over the course of learning. **A)** The ratio of within- and between-class variance was computed in the maximally discriminative subspace for object ‘identity’ (*κ* = 15 classes, inset top left) and object ‘novelty’ (2 classes, inset bottom left). On the right, the average values 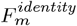 and 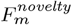, over observers and identity-selective parcels and for successive batches *m* are shown in terms of z-score values 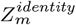 (solid blue line and shading, mean ± S.E.M.) and 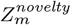 (solid red line and shading, mean ± S.E.M). While 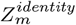 decreases slightly over time (*p* < 0.05), 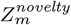 increases considerably (*p* < 0.001), especially initially. **B)** To reveal the respective contribution of within- and between-class variances, the values for 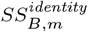 (dashed blue), 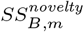 (dashed red), 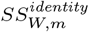 (solid blue), and 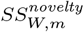 (solid red) are shown relative to the corresponding shuffled values (mean± S.E.M.). While 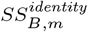 decreased slightly over time (*p* < 0.05), 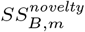 increased considerably (*p* < 0.001). **C)** Pairwise correlation between ‘slope’ parameters 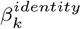 and 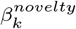, and identification accuracy *a*_*k*_, over parcels *w*. Parameters 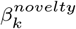 was only weakly correlated with parameter 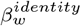 (*ρ* = 0.298, *p* < 0.001) (left) and accuracy *a* (*ρ* = − 0.22, *p* < 0.05) (middle), whereas 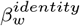 and *a*_*k*_ were strongly correlated (*ρ* = − 0.74, *p* < 0.001)(right).

To distinguish whether it was the within-or the between-class variance that changed over time, the development of within- and between-class variances is shown separately as well. Specifically, the values of 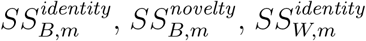, and 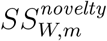 are illustrated in **Fig. 4B**, relative to the corresponding shuffled values. The results show that it was between class-variance 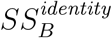 and 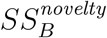 that changed significantly over successive batches *m* (*p* < 0.05 and *p* < 0.001, respectively), whereas with-class variance 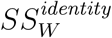 and 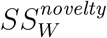 remained essentially unchanged (*p* = 0.75 and *p* = 0.58, respectively).

In other words, over the course of learning, the representations of different recurrent objects tend to become slightly more similar to each other, whereas the representation of non-recurrent objects becomes more dissimilar to that of recurrent objects.

To summarize the change observed in particular parcels, we estimated ‘slope’ pa-rameters 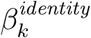 and 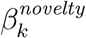 by fitting variance ratios 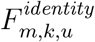 and 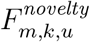 by linear mixed-models. Over all identity-selective parcels, 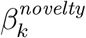 varied more than 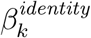, as measured by standard deviation *σ*(*β*^*identity*^) = 0.015 and *σ*(*β*^*novelty*^) = 0.097.

The joint distribution of slopes 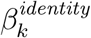 and 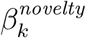 over parcels identity-selective parcels *k* is illustrated in **Fig. 4C**. Identity slope was correlated to novelty slope (*ρ* = 0.298, *p* < 0.001), and both identity slope and novelty slope were anti-correlated with identity selectivity (*ρ* = −0.22, *p* < 0.05 and *ρ* = −0.74, *p* < 0.001, respectively).

The degree to which parcels *k* express the overall novelty trend, as measured by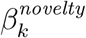, differs between regions of the brain. Interestingly, the most pronounced representation of novelty was found in parietal and frontal areas, rather than in occipital areas (**Fig. 5.A, Table A.1**,**Fig. S4**).

**Figure 5:**
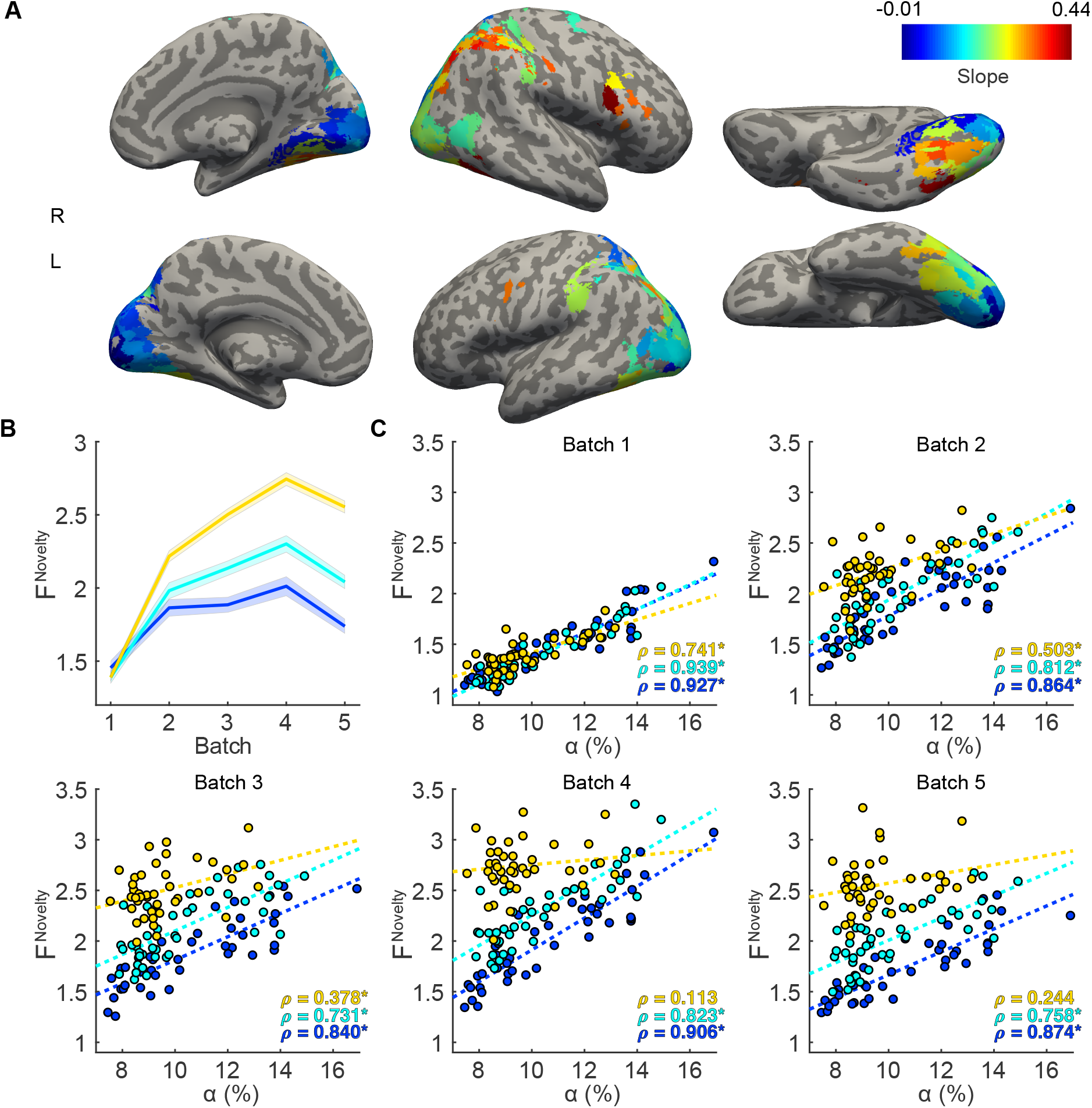
Neural representation of ‘novelty’ (non-recurring objects). **A)** Distribution of rate parameter *β*^*novelty*^ over identity-selective parcels. Parameter 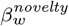 is the slope or rate of increase of 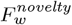 of parcel *w*, estimated by a linear mixed-model. **B)** Development of *F*^*novelty*^ in upper, middle, and lower tercile of parcels (mean ± SEM), defined in terms of *β*^*novelty*^. **C)** Correlation between 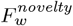 and accuracy *α*_*w*_ for different batches and different terciles (distinguished by color). Regression lines (dashed) and correlation coefficients *ρ* are also shown (*indicates *p* < 0.05).

To examine changes in the representation of ‘novelty’ (non-recurring objects) more closely, we divided the parcels into terciles in terms of their rate parameter *β*^*novelty*^ (high, medium, and low rate) (**Fig. 5.B**). The results differed between batches and terciles. In early batches, *F* ^*novelty*^ and classification accuracy *α* were correlated in all terciles. In other words, at the beginning of learning, the representation of non-recurring objects (indexed by *F* ^*novelty*^) and recurring objects (indexed by *α*) were linked in all parcels. This correlation waned in successively later batches, but only in the upper tercile of *β*^*novelty*^. Accordingly, it seems that the representation of non-recurrent objects became successively more independent of the representation of recurrent objects, at least when the latter representation was particularly pronounced (upper tercile of *β*^*novelty*^). In such parcels, the respective representations of non-recurrent and recurrent objects appeared to become independent of each other.

Somewhat contrary to our expectations, identity-related representations became evident already during first experimental session, not just in the last session after objects had become wholly familiar to observers. However, these early signs for identity-related representations were consistent with the behavioral observations that most objects become familiar already during the first session (Kakaei et al., 2021). As mentioned above, spatiotemporal activity may have reflected invariant relationships between multiple object features. Whatever the nature of this encoding, this was clearly a general-purpose representation capable of distinguishing arbitrary complex objects from the start. The *quality* of this representation remained largely stable over the course of learning, apart from growing slightly less pronounced over time. In other words, our results are consistent with the consolidation of a pre-existing representation of complex objects, not with the gradual formation of a novel representation.

In fact, the most pronounced change over the course of learning was observed in the representation of non-recurrent objects. For the representational distance between non-recurrent (‘novel’) and recurrent (‘familiar’) objects increased substantially over the course of learning. In contrast, the introspective experience and behavioral response of observers changed with respect to both recurrent and non-recurrent objects: the former came to be recognized as ‘familiar’ and the latter (by default) as ‘novel’.

### 3.3 Quality of identity and novelty representations

To investigate neural representations and their changes with learning more closely, we compared geometrical properties of non-recurrent (novel) and recurrent (familiar) object representations in a fixed discriminate space and over successive ‘runs’ (each comprising 200 trials, see Methods for details). By using a fixed space, we hoped to improve the spatial characterization of representational geometry. By analyzing successive runs, we hoped to improve the temporal resolution of the analysis. Specifically, we analyzed representational geometry in terms of individual response amplitude, the distance between response pairs, and variance of distance between response pairs, separately for each of the 18 runs of the experiment, which were performed in three sessions. All distances were residual distances and corrected to remove temporal auto-correlations (**Supplementary Fig. S1**; see Methods for details).

Results are summarized in **Fig. 6**, separately for the 124 parcels identified previously as identity-selective and the 634 remaining parcels. The analyzed quantities – average pattern amplitudes *A*, average pattern distances *D*, and variance ratios *SS* – are explained and illustrated schematically in **Fig. 6A**. In identity-selective parcels (**Fig. 6B**), response amplitudes *A*_*fam*_ to recurring patterns decreased during the first session (runs 1 to 6, *p* < 0.05), but not in the second and third session (runs 7 to 12, runs 13 to 18, *p* > 0.5). Response amplitudes *A*_*nov*_ to non-recurring patterns show no significant trend in any session (*p* > 0.36). Response distance *D*_*diff*_ between recurring patterns declined similarly during the first session (*p* < 0.05), but not during subsequent sessions (*p* > 0.6). (**Fig. 6C**). Response distance *D*_*same*_ did not change during the sessions (*p* > 0.15). Response distances *D*_*nov*_ between non-recurring patterns declined disproportionately during the first session (*p* < 0.05), but increased during the third session (*p* < 0.05). Response distances *D*_*novfam*_, on the other hand, did not change significantly over sessions (*p* > 0.13). Variances *SS*_*same*_ and *SS*_*diff*_ for recurring objects showed different trends: while *SS*_*diff*_ declined over all sessions (*p* < 0.005), *SS*_*same*_ remained unchanged (*p* > 0.05). For non-recurring objects, variance *SS*_*novfam*_ increased over the first session (*p* < 0.005), but declined during the third session (*p* < 0.05). Variance *SS*_*nov*_, on the other hand, decreased over the first session (*p* < 0.05) and afterward remained unchanged. Interestingly, the lowest distances and variances were observed for non-recurring patterns (*D*_*nov*_ and *SS*_*nov*_), consistent with poor representational quality.

**Figure 6:**
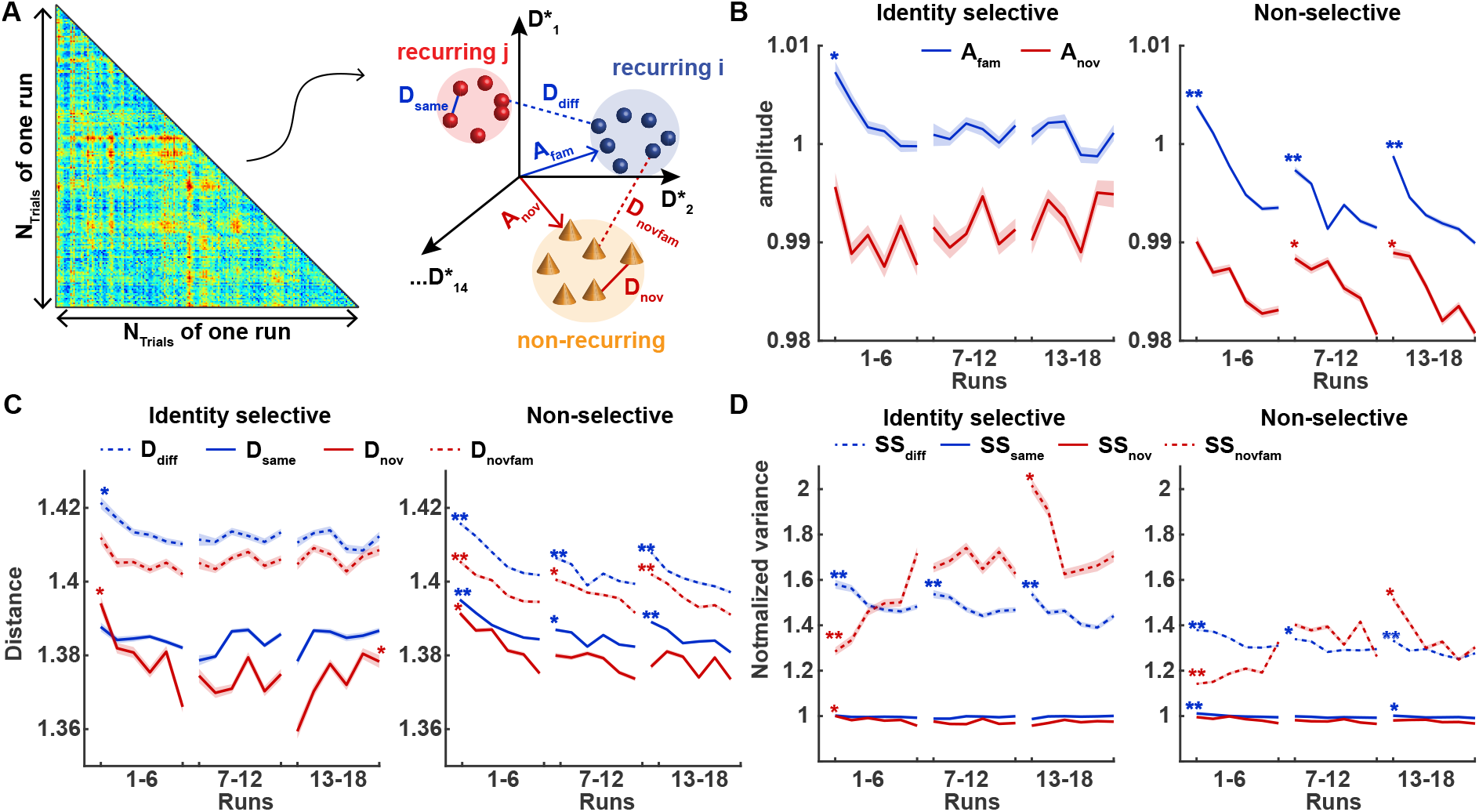
Quality of identity and novelty representation over the course of learning. **A)** For each run with *N*_*trials*_, we computed all individual response amplitudes *A* and pairwise distances *d* in the maximally discriminating space and obtained average amplitudes *A*_*fam*_ and *A*_*fam*_ (for recurrent and non-recurrent objects, respectively) and average distances *D*_*same*_ and *D*^*diff*^ (for same and different recurrent objects, respectively), as well as average distances *D*_*nov*_ and *D*_*novfam*_ (for non- recurrent objects and between recurrent and non-recurrent objects, respectively). **B)** Normalized amplitude *A*_*nov*_ (red) and *A*_*fam*_ (blue), over 18 runs, grouped into three sessions, for identity-selective and non-selective parcels. **C)** Normalized pairwise distance *D*_*same*_ (solid blue), *D*_*diff*_ (dashed blue), *D*_*nov*_ (solid red), and *D*_*novfam*_ (dashed red)), over runs and sessions, for both sets of parcels. **D)** Variances ratios *SS*_*same*_ (solid blue), *SS*_*diff*_ (dashed blue), *SS*_*nov*_ (solid red), and *SS*_*novfam*_ (dashed red)), over runs and sessions, for identity-selective and non-selective parcels. Stars indicate a significant linear trend during a session (see text).

As shown in Supplementary **Figure S5**, the decreasing separation between *SS*_*same*_ and *SS*_*diff*_ has the variance fraction *R*^*identity*^ ∝ *SS*_*diff*_ */SS*_*same*_ mirror the trend observed for *F* ^*identity*^ in the earlier batch analysis (**Fig. 4. A**), whereas the increasing separation between *SS*_*nov*_ and *SS*_*novfam*_ during the first session (and decreasing separation during the third session) makes variance fraction *R*^*novelty*^ ∝ *SS*_*novfam*_ */* (*SS*_*fam*_ + *SS*_*nov*_) mirror the trend observed earlier for *F* ^*novelty*^.

In non-identity-selective parcels, amplitudes, distances and variances declined progressively in every session, except for the variance between non-recurring and recurring objects *SS*_*novfam*_ which increased initially and subsequently declined (**Fig. 6B-D**). Note that amplitudes and distances were generally *smaller* than in identity-selective parcels, consistent with lower representational quality. An exception was the distance *D*_*nov*_ between non-recurring objects, which was larger in non-identity-selective parcels. The separation between *SS*_*same*_ and *SS*_*diff*_, as well as that between *SS*_*nov*_ and *SS*_*novfam*_, was considerably smaller than in identity-selective parcels, again consistent with lower representational quality.

Several general observations are noteworthy. Firstly, in all parcels, amplitudes and distances are larger between classes than within classes of recurring patterns, presumably because responses to similar stimuli are more similar than responses to dissimilar stimuli. Secondly, in all parcels, amplitudes and distances are larger for recurring than for non-recurring patterns, presumably because the representational space S was optimized for the former, not for the latter. Thirdly, in non-identity-selective parcels, amplitudes and distances decrease progressively within and between sessions, presumably because neuronal response habituates and/or hemodynamic impact declines. However, in identity-selective parcels, this decline was arrested and amplitudes and distances stabilized during the first session. Fourthly, between-class distances were larger in identity-selective than in non-identity-selective parcels (while within-class distances were comparable), as expected for a more informative representation. Lastly, the situation appeared somewhat different with regard to the respective representation of recurring and non-recurring objects. Although between-class distances were once again larger, within-class distances for non-recurring objects were smaller, when comparing identity-selective and non-identity-selective parcels. In fact, the initial decline (and later partial recovery) of within-class distances for non-recurrent objects constituted the most prominent neural correlate of object recognition learning that was evident in our data.

### 3.4 Stability of identity and novelty representations

The analyses of the previous section focused on the *quality* of representation, that is, on relative distances within and between classes of response patterns. However, as representations were specified in the same discriminative subspace S over all runs, it became possible to additionally assess the *stability* of representation over the course of learning. In principle, quality and stability are independent and could conceivably develop differently over the course of learning. To assess relative stability between successive runs and absolute stability over all runs, we computed centroids of response patterns for each run and established the movement of centroids from one run to the next as well as over all runs. As these comparisons pertained to centroid-to-centroid distances (rather than exemplar-to-exemplar distances) they could not be corrected for temporal auto-correlations.

Run-specific centroids *C*_*r−*1_ and *C*_*r*_ and overall centroid *C*_*all*_ are illustrated schematically in **Fig. 7A**. Absolute centroid-to-centroid distances *D*(*r*) were computed between *C*_*r*_ and *C*_*all*_ and relative centroid-to-centroid distances Δ*D*(*r*) between centroids *C*_*r−*1_ and *C*_*r*_ of successive runs. The results were quite unexpected and are summarized in **Fig. 7B**. For both recurring and non-recurring objects, average absolute distances *D*_*same*_ (*r*) and *D*_*nov*_ (*r*) diminished during the first session (runs 1 to 6, *p* < 0.005), but remained stable during the second and third sessions (runs 7 to 12, and 13 to 18, *p >* 0.2). Notably, absolute distances *D*_*nov*_ (*r*) of novel objects decreased to a much lower average level, consistent with a more compact representation. Relative distances Δ*D*_*same*_ (*r*) and *D*_*nov*_ (*r*) between successive runs declined during the first session (runs 1 to 6, *p* < 0.05), remained stable during the second session (runs 7 to 12, *p >* 0.2), only to decline once again the last session (13 to 18, *p* < 0.005 for recurring and *p* < 0.05 for non-recurring objects). Astonishingly, the relative distances between successive runs were *larger* than the absolute distances to the overall centroid. This was particularly true for recurring objects and reveals a remarkable degree of instability in their representation.

**Figure 7:**
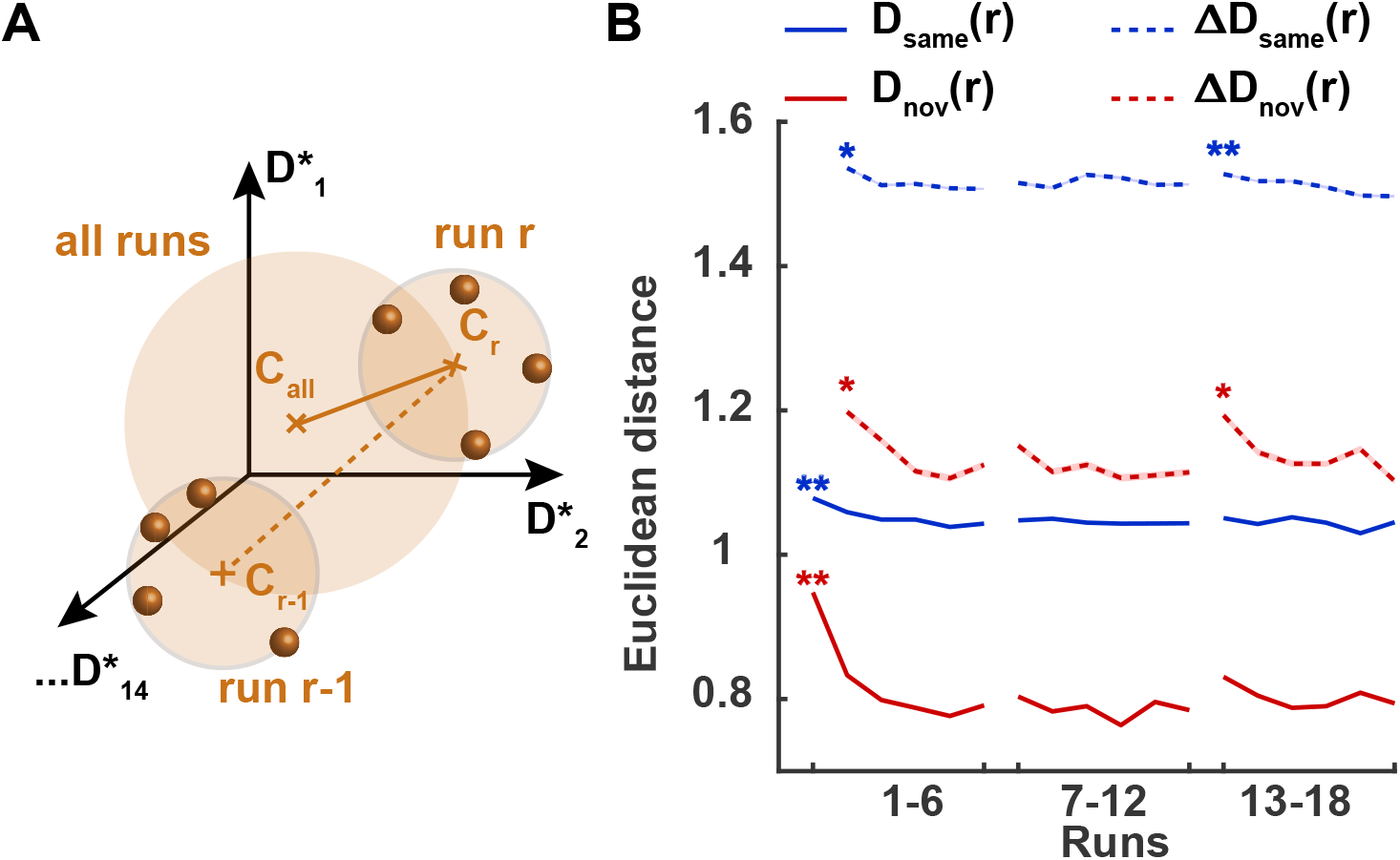
Stability of identity and novelty representation over the course of learning. **A)** For each recurring and all non-recurring objects, we calculated a grand centroid **C**_**all**_ over all the runs and calculated a centroid **C**_**r**_ over run *r*. Then, Euclidean distance *D*(*r*) (solid curves) between the grand centroid and the centroid of an individual run was calculated. Additionally, we calculated distance Δ*D*(*r*) (dashed curves) between the relevant centroids from successive runs. **B)** Centroid-to-centroid distances averaged over identity-selective parcels and participants (*±S*.*E*.*M*), are shown for recurring objects (with *D*_*same*_ (*r*) in solid blue and *ΔD*_*same*_ (*r*) in dashed blue) and non-recurring objects (with *D*_*nov*_ (*r*) in solid red and Δ*D*_*nov*_ (*r*) in dashed red). Stars indicate a significant linear trend during a session (see text).

In summary, we can say the following about relative and absolute movement over the course of learning. Firstly, movement is substantially smaller for the representation of non-recurring objects, consistent with their more compact representation (see above). Secondly, relative movement between successive sessions is substantially *larger* than absolute movement over all sessions, revealing a remarkable instability of representation, especially for recurring objects. It would appear that representation of recurring objects changes continuously during the course of learning, even though the quality of this representation remains essentially the same.

## 4 Discussion

We studied the cortical representation of synthetic visual objects over multiple days of experience, during which time initially unfamiliar objects gradually became familiar. Relying on ‘representational similarity analysis’ (RSA), we established distances between spatiotemporal BOLD responses to exemplars of different *recurring* objects, as well as to exemplars of *non-recurring* objects. To quantify the representation of object *identity*, we compared distances between the same and different recurring objects. To assess the representation of object *novelty*, we compared distances between recurring and non-recurring objects. Object identity was represented from the first day in both the ventral and dorsal pathways. With additional experience, the quality of representation barely declined, even though the geometry of representation continued to change. The respective representations of recurring and non-recurring (novel) objects became increasingly distinct and the latter became increasingly marginalized.

Our approach to RSA differed from previous work in some respects. We analyzed comparatively high-dimensional *spatiotemporal* patterns of BOLD activity (200 voxels × 9 seconds, or O(2000) dimensions) in non-overlapping gray matter volumes (758 functional subdivisions of 90 anatomical regions, averaging 1.7 cm^3^, Dornas and Braun 2018), rather than lower-dimensional *spatial* activity patterns in overlapping searchlight volumes (*O*(50) voxels or dimensions, covering 0.25 to 1.0 cm^3^; Kriegeskorte et al. 2006). For each gray matter volume and its high-dimensional activity space, we determined the subspace that optimally distinguished responses to recurring objects (*i*.*e*., a 14-dimensional space distinguishing 15 object classes), using multi-class linear discriminant analysis (‘direct linear discriminant analysis’, DLDA; Yu and Yang 2001), rather than relying on pairwise discriminability or one-versus-all discriminability (*e*.*g*., Liu et al., 2009; Hung et al., 2005). With these modifications, RSA revealed representational geometry at the level of object exemplars or subclasses, as well as gradual changes in this geometry with additional experience and familiarity.

### Representation of object identity

To facilitate fine-grained analysis of representational geometry, we developed synthetic shapes for which visual expertise was acquired comparatively slowly (Kakaei et al., 2021). While these complex and three-dimensional shapes were highly characteristic and distinguishable, they were shown from variable points of view while rotating about variable axes (for one complete turn). Accordingly, in order to categorize an object as ‘familiar’, observers had to recognize its appearance from different points of view. In effect, the different ways of presenting a recurring object constituted different ‘exemplars’ of a particular ‘object’, with each ‘object’ forming a subclass within the superclass of our synthetic shapes.

The selectivity of cortical parcels for object identity was assessed in terms of crossvalidated ‘classification accuracy’ and prevalence analysis served to combine the results from all observers. An alternative measure, which yielded essentially identical results, was Anderson’s ‘variance ratio’ (Anderson, 2001). This compares exemplars of *different* objects and exemplars of the *same* object in terms of pairwise distances between spatiotemporal responses. A variance ratio above unity implies that the spatiotemporal responses to different objects are linearly discriminable and therefore a representation of object identity. As exemplars of the same object are shown from varying points of view, any such representation is view-invariant by definition. Of course, the presence of a view-invariant representation does not exclude the possibility that a view-specific representation is also present, perhaps in some other subspace of the high-dimensional response space.

The 124 of 758 parcels that were identified as ‘identity-selective’ on this basis were situated mostly in the ventral occipitotemporal cortex, but some parcels were located also in the parietal or frontal cortex, as summarized in **Fig. 8A**. The degree of selectivity exhibited a clear gradient, being stronger at the posterior pole and becoming progressively weaker in more anterior and more dorsal regions. The cortical distribution of the observed representation is entirely consistent with previous findings that multivariate activity distinguishing different exemplars of a particular class of objects (*e*.*g*., faces) is present in the ventral and lateral occipital cortex, on the fusiform gyrus, and in ventral temporal cortex (Eger et al., 2008; Brants et al., 2016; Visconti di Oleggio Castello et al., 2021).

**Figure 8:**
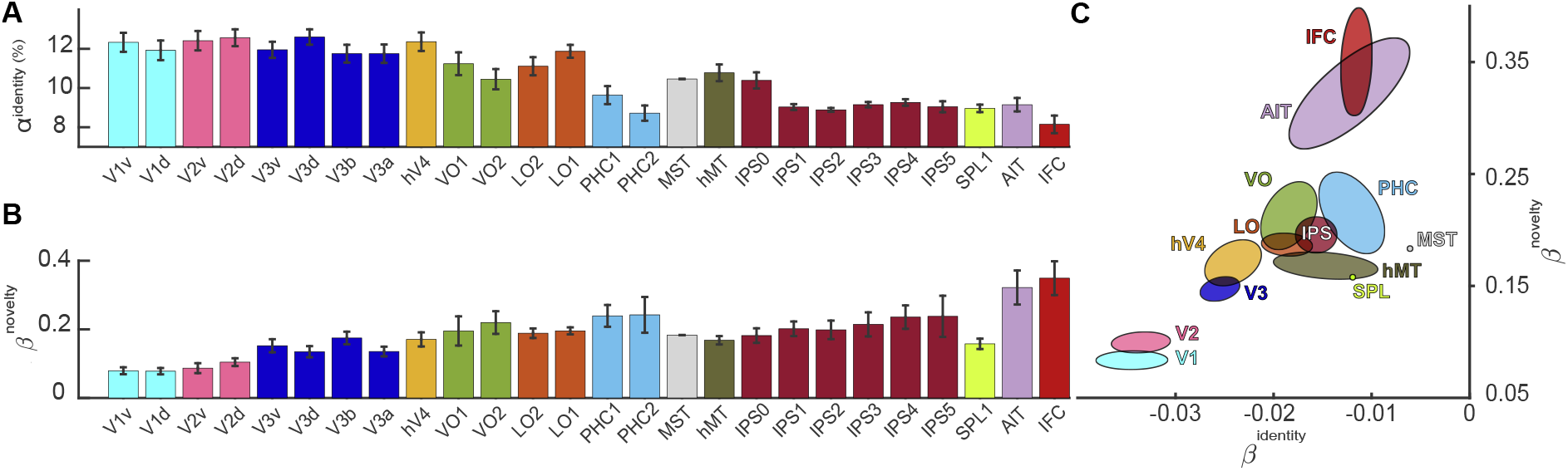
Identity and novelty representations in 26 topographical regions **A)** Identity representation as indexed by classification accuracy *α*^*identity*^ (mean± S.E.M.). Posterior regions (V1-hV4, VO, LO) exhibit higher accuracy than more anterior or more dorsal regions (IPS, AIT, IFC). **B)** Novelty representation as indexed by rate *β*^*novelty*^ of novelty gain (mean ±S.E.M.). More anterior or more dorsal regions (IPS, AIT, IFC) exhibit a higher slope parameter than posterior visual cortex (V1-hV4). **C)** Comparison of identity and novelty representations as indexed by rate *β*^*identity*^ of identity loss (negative values) and rate *β*^*novelty*^ of novelty gain. Groups of regions are distinguished by color, with ellipsoids indicating mean and standard error. Note the negative correlation between novelty *gain* and identity *loss*. List of abbreviations: visual cortex (V1, V2, V3, hV4), ventral occipital cortex (VO1, VO2), lateral occipital cortex (LO1, LO2), parahippocampal cortex and fusiform gyrus (PHC), medial temporal areas (hMT, MST), intraparietal sulcus (IPS), superior parietal lobule (SPL), anterior inferior temporal cortex (AIT), inferior frontal cortex (IFC).

A small discrepancy is that we observed identity-selectivity also in the primary visual cortex (calcarine sulcus), where multivariate activity is typically expected to be selective for basic visual features (orientation, spatial frequency, direction of movement, and so on) rather than object categories (Haxby et al., 2001; Grill-Spector and Weiner, 2014). Of course, identity-selectivity of a cortical cluster may well be due to (feedback) projections from other cortical clusters, rather than arising locally with the cluster itself. Patterns of BOLD activity are particularly susceptible to such ‘functional correlations’. Distinguishing between clusters in which identity selectivity originates and clusters that merely echo such selectivity would require an analysis of statistical symmetries such as transfer entropy or Granger causality and is beyond the scope of the present study.

The observation that identity selectivity is less pronounced in more anterior than in more posterior parts of the ventral temporal cortex is consistent with established views. In general, it is hypothesized that progressively ‘higher’ levels of visual processing represent progressively ‘larger’ visual sets, beginning with image features, and widening gradually to object features, object exemplars, object categories, and finally to supercategories such as animate or inanimate objects, or objects and landscapes (Grill-Spector and Weiner, 2014). Accordingly, the discriminability of exemplars within a category is expected to diminish at more anterior locations, which correspond to ‘higher’ levels of visual processing (Eger et al., 2008; Grill-Spector and Weiner, 2014). Moreover, it has been hypothesized that the spatial scale of neural representations increases with the level of abstraction, in the sense that exemplars are represented at smaller scales than categories (Grill-Spector and Weiner, 2014). Thus, if this trend is exacerbated in the more anterior parts of the ventral pathway, exemplar representations may become progressively less discriminable at the spatial resolution of BOLD signals.

Our results demonstrate identity selectivity not just in the ventral occipitotemporal cortex, but also in frontoparietal regions that are typically associated with the dorsal visual pathway and the right frontoparietal “attention network”. This is consistent with previous findings on the presence of object- and/or face-selective representations in dorsal areas (Poirier et al., 2006; Konen and Kastner, 2008; Jeong and Xu, 2016; Freud et al., 2017; Visconti di Oleggio Castello et al., 2021). However, while identity-selectivity in parietal clusters might have arisen locally with the parietal cortex itself, it might also have been due to (feedforward) projections from other cortical clusters. Particularly the clusters associated with the “attention network” are often found to express functional correlations with ventral visual areas. In the context of the present study, it is conceivable that such functional correlations could propagate identity-selectivity through this “attention network”.

### Representation of object novelty

We also investigated the representation of ‘novel’ object shapes that were encountered exactly once (*i*.*e*., did not recur). Note that ‘novelty’ here does not necessarily imply a ‘surprise’ in the sense of a visually salient violation of expectations (Uddin, 2015). Specifically, in the optimally discriminating subspace for recurring objects, we compared exemplars of *non-recurring* and *recurring* objects in terms of pairwise distances between spatiotemporal responses. Between-group distances larger than within-group distances (*i*.*e*., variance ratio above unity) implied that non-recurrent and recurrent objects were linearly discriminable and that the ‘novelty’ of object shape was represented in the BOLD responses.

All 124 ‘identity-selective’ parcels were also ‘novelty-selective’ to some degree, as summarized in **Fig. 8B**, which suggests that the observed representation is general and pertains to all shapes, rather than being specific and pertaining exclusively to particular shapes. Note that the approximately 300 non-recurring objects constituted an intrinsically more variegated set than the 15 recurring objects and that there was no systematic difference between non-recurring and recurring shapes. Interestingly, novelty-selectivity was more pronounced in more anterior and more dorsal frontal, parietal, and temporal areas, than in posterior occipital areas. The relation between identity- and novelty-selectivity in different parcels will be discussed further below, in the context of changes with learning.

### Representational changes with learning

To assess representational changes with learning, we pursued two complementary approaches. In the first approach, we divided our observations from eighteen runs (performed in three sessions) into five successive ‘batches’, favouring cross-validated statistics over temporal resolution and a stable discriminative space. In the second approach, we analyzed individual runs, sacrificing cross-validation to gain temporal resolution and a stable discriminative space.

Identity representations were maximally differentiated already during the first batch and after the first few runs. This initial identity representation was most pronounced in the core object processing areas, including early visual cortex and ventral occipitotemporal cortex. We hypothesize that pre-existing representations based on life-long experience were sufficient to rapidly provide a view-independent representation of synthetic shapes, which we had designed to be highly characteristic and discriminable. In contrast, there was no initial discriminability of recurring and non-recurring objects and thus no initial representation of novelty. This was expected because there was no systematic difference between recurring and non-recurring objects, so that observers could not have distinguished them at the start of the experiment.

While the representation of object identity remained pronounced, it declined slightly but significantly over the course of the experiment. Some decline in BOLD activity is not untypical for learning studies over multiple days and is commonly ascribed to repetition suppression, sparsification of responses, and/or diminishing attention or effort (*e*.*g*., Poldrack, 2000).

A previous study of visual expertise for synthetic shapes reported a gradual enhancement of subclass representation in object-selective areas, specifically in lateral occipital cortex (Brants et al., 2016). The reason for this difference may have been the behavioural task. Whereas Brants and colleagues required the discrimination of two subclasses of objects that were barely discriminable, we merely asked for the discrimination of superclasses (familiar or novel?) of objects that were highly discriminable. In short, the paradigm of Brants and colleagues emphasised perceptual load, whereas our paradigm put more stress on memory load.

We hypothesize that observers may have formed a mental category of ‘familiar’ objects over the course of the experiment, which would have been sufficient for the behavioral task, but would not have required detailed information on object identity. If attentional feedback from right frontoparietal regions to early visual areas (Zanto et al., 2010, 2011) increasingly emphasized this categorical information at the expense of detailed shape information, this could explain the observed decrease in identity representation.

The linear discriminability of recurring and non-recurring objects improved substantially over the course of the experiment, revealing an enhanced representation of novelty. The gradual time-course of this improvement was similar in both analyses (batch-by-batch and run-by-run). In contrast, the behavioural performance of observers in distinguished recurring and non-recurring objects approached ceiling already during the first ‘batch’ (after three runs). Accordingly, the observed representational changes do not appear to correspond closely to improvement of behavioural performance.

The general pattern of the observed representational changes is illustrated in **Fig. 9**, where two-dimensional positions approximately preserve relative pairwise distances in the 14-dimensional response space. While response distances to recurrent objects decrease over the course of the experiment, different recurrent objects remain linearly discriminable (**Fig. 9A**). In contrast, response distances between recurrent and non-recurrent objects increase over the course of the experiment (**Fig. 9B**). In fact, the representation of non-recurrent objects becomes increasingly marginalized in the response space, even though non-recurrent objects constitute the more variegated set. We hypothesize that the representation of the fifteen recurrent objects, which observers are tasked to memorize, expands during learning and ‘fills’ the available response space. In doing so, this representation progressively displaces the representation of non-recurring objects, which observer observers are not trying to memorize.

**Figure 9:**
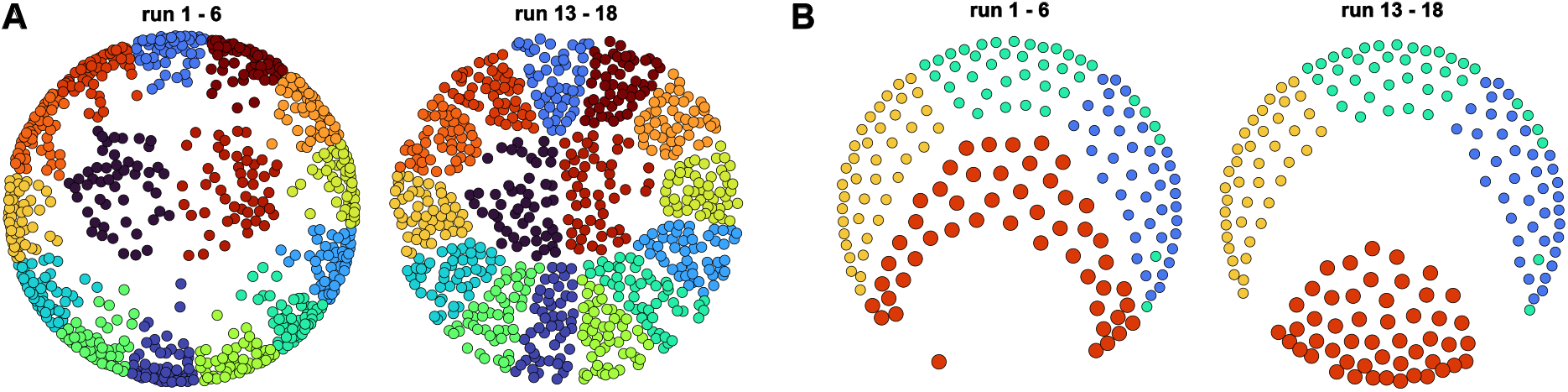
Changes in the geometry of identity and novelty representations, visualized by multi-dimensional scaling. Here, two-dimensional position reflects relative pairwise distances in a 14-dimensional discriminative space. **A)** Randomly selected exemplars of fifteen recurring objects (distinguished by color) in the first session (left) and the last session (right). **B)** Randomly selected exemplars of non-recurring objects (large red circles) in the first session (left) and the last session (right), relative to exemplars of three recurring objects (distinguished by color) from either session.

In addition to relative changes in representational geometry indexed by linear discriminability, we also observed absolute changes in representational geometry, indexed by distances between response centroids in successive runs. Suprisingly, response centroids moved considerably between successive runs, revealing substantial ongoing changes with continued experience (see **Fig. 7**). Simultaneously, response centroids remained in the proximity of their respective average locations, safeguarding the representation of identity. In other words, the movement of response centroids was comparable to random jumps on the circumference of a circle, maintaining a constant distance from its center. We conclude that the comparative stability in the *quality* of representations concealed a massive instability in the *geometry* of representations, in that the latter continued to be modified with further visual experience.

The rate of change in the quality of identity and novelty representations differed substantially between cortical regions, as summarized in **Fig. 8C**. Intriguingly, over all levels of the shape processing hierarchy, the rates of novelty *gain* and identity *loss* changed systematically (and inversely): in early visual areas (V1, V2, V3, hV4), identity declined rapidly, whereas novelty grew slowly. At the opposite end, in inferior frontal cortex (IFC) and anterior ventral temporal cortex (AIT), identity declined slowly, but novelty grew rapidly. In higher visual cortex (VO, LO), both rates were intermediate. Some areas deviated somewhat from this trend: dorsal and parietal areas (hMT, MST, IPS, SPL), as well as fusiform and parahippocampal cortex (PHC), also lost identity slowly, but gained novelty at an intermediate pace.

These results suggest a partial dissociation between the representations of identity and of novelty. This conclusion is further strengthened by comparing the rate of novelty gain to the absolute level of identity-selectivity (see **Fig. 5**): some parcels with rapid novelty gain were not particularly identity-selective (PHC, IPS, AIT, IFC). Accordingly, it seems clear that the development of identity-and novelty-selectivity while visual expertise is accumulated and consolidated is convergent in some cortical areas, but divergent in others.

## Conclusion

Recent studies of visual expertise have highlighted the roles of three pathways or networks (Kravitz et al., 2011, 2013), an occipito-temporal pathway (“ventral pathway”), an occipito-parietal pathway (“dorsal pathway”), and a right frontoparietal network (“attention system”). Several studies linked behavioural performance to enhanced activity and/or representation in the frontoparietal network (Poirier et al., 2006; Visconti di Oleggio Castello et al., 2021; Duyck et al., 2021), as well as in the more anterior parts of the occipito-temporal pathway and the more dorsal parts of the occipito-parietal pathway (Christophel et al., 2017). The development of novelty representations that we observed in these regions further corroborates these earlier findings.

The most robust representations combining object identity and novelty were located in ventral occipitotemporal cortex, at the intermediate levels of the shape processing hierarchy (Perry and Fallah, 2014; Grill-Spector and Weiner, 2014). That view-independent representations of object exemplars should be present in these particular cortical regions was entirely expected. That these representations were present already from the start, we attribute to the highly discriminable and characteristic nature of our synthetic stimuli. We surmise that less discriminable shapes might have produced a different outcome (Brants et al., 2016).

Perhaps the most interesting result of the present study was the continuing change in the representation of recurring objects with further experience, which seemed to progressively ‘fill’ the discriminative dimensions of the response space. In contrast, the representation of non-recurring objects seemed to become progressively marginalized. Thus, our results demonstrate that the acquisition of visual expertise differentiates between the representation of similar synthetic objects, with familiar objects being afforded progressively more space and unfamiliar objects being afforded less. Further work will be required to more fully understand the precise influence of the behavioural task, which in the present study merely required identifying objects as “familiar”. Conceivably, representational geometry for the same stimulus set might have developed differently in the context of other behavioral tasks, for example, a requirement to identify specific objects “by name”.

## Data and code availability

Direct linear discriminant analysis and prevalence inference is available on github.com/cognitive-biology/DLDA. MR data will be made available upon request.

## Credit authorship contribution statement

**E. Kakaei:** Conceptualization, data curation, formal analysis, visualization, writing of original draft. **J. Braun:** Conceptualization, linear algebra, formal analysis, supervision, reviewing & editing.

## Acknowledgments

We thank Claus Tempelmann, Martin Kanowski and Denise Scheermann at the Magnetic Resonance Imaging Laboratory of the Department of Neurology of Otto-von-Guericke University, Magdeburg. We also thank Stepan Aleshin for helpful discussions and constructive comments. This study was funded by the federal state Saxony-Anhalt and the European Structural and Investment Funds (ESF, 2014-2020), project number ZS/2016/08/80645, as part of doctoral program ABINEP (Analysis, Imaging and Modelling of Neuronal Processes).

## A Appendices

**Table A.1:**
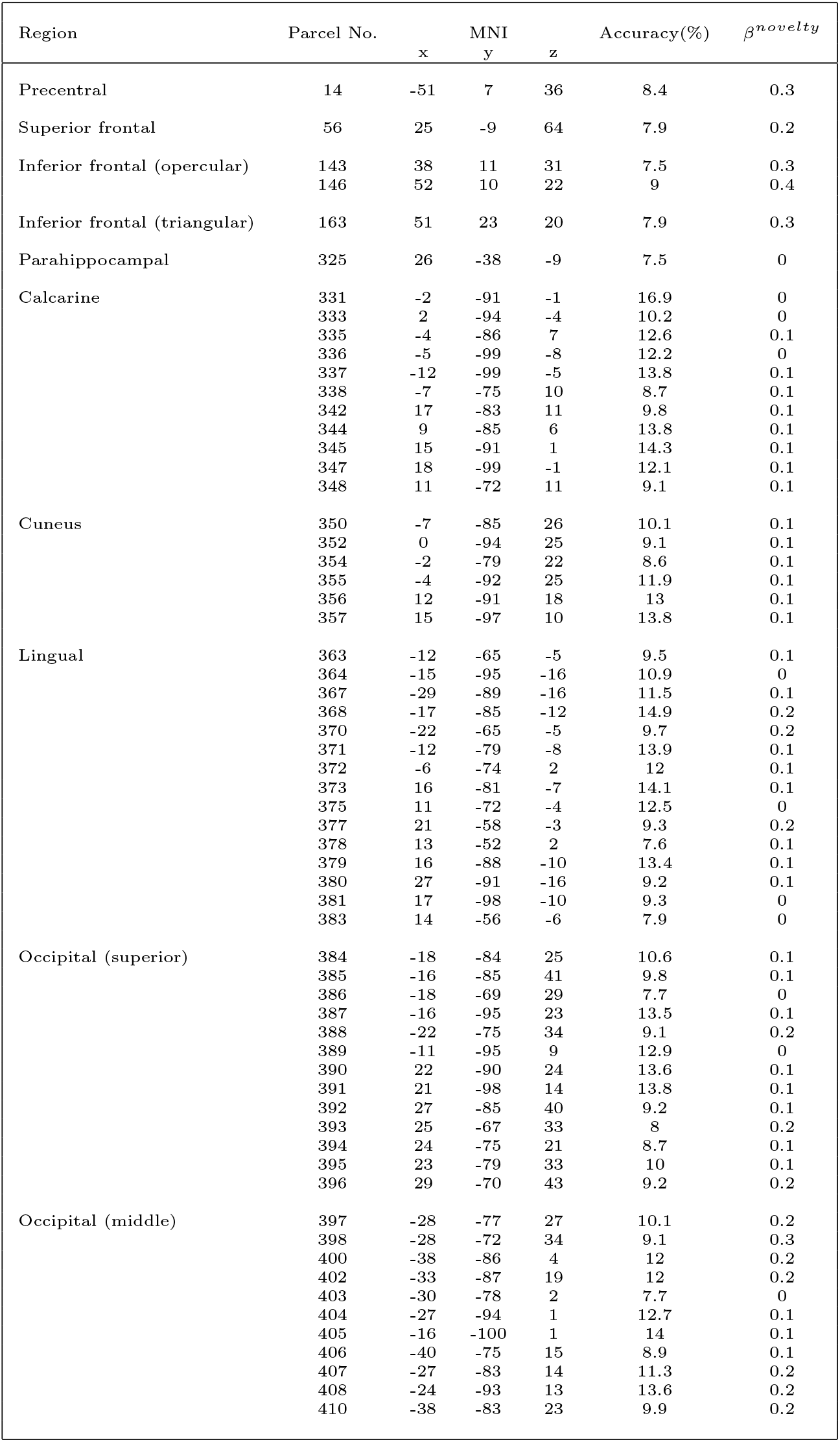
List of identity selective parcels and their anatomical region. Numerical parcel ID, geometrical centroid x/y/z in MNI, mean cross-validation accuracy, and novelty gain parameter *β*^*novelty*^.

**Table A.2:** List of identity selective parcels and their anatomical regions: numerical parcel ID, geometrical centroid x/y/z in MNI, mean cross-validation accuracy, and novelty gain parameter *β*^*novelty*^.

| Region | Parcel No. | MNI | | | Accuracy(%) | $\beta^{novelty}$ |
| --- | --- | --- | --- | --- | --- | --- |
|  |  | x | y | z |  |  |
|  | 412 | -46 | -77 | 5 | 10.1 | 0.1 |
|  | 413 | 33 | -88 | 7 | 13.2 | 0.2 |
|  | 414 | 33 | -96 | 4 | 10.7 | 0.2 |
|  | 415 | 39 | -81 | 14 | 10.6 | 0.2 |
|  | 416 | 44 | -78 | 5 | 10.9 | 0.2 |
|  | 418 | 32 | -86 | 23 | 11.9 | 0.2 |
|  | 420 | 34 | -69 | 32 | 8.5 | 0.3 |
|  | 421 | 32 | -76 | 27 | 10 | 0.3 |
| Occipital (inferior) | 423 | -50 | -68 | -14 | 9.4 | 0.2 |
|  | 424 | -31 | -83 | -8 | 12.6 | 0.2 |
|  | 425 | -22 | -95 | -9 | 12.3 | 0.1 |
|  | 426 | -42 | -73 | -8 | 11.6 | 0.2 |
|  | 428 | 36 | -85 | -7 | 13.2 | 0.2 |
|  | 430 | 42 | -73 | -9 | 11.4 | 0.3 |
| Fusiform | 432 | -27 | -71 | -11 | 12.2 | 0.2 |
|  | 435 | -33 | -77 | -17 | 12.4 | 0.1 |
|  | 436 | -31 | -53 | -13 | 8.8 | 0.3 |
|  | 438 | -41 | -56 | -17 | 8.9 | 0.3 |
|  | 440 | -36 | -63 | -16 | 9.6 | 0.3 |
|  | 442 | 28 | -74 | -11 | 12.8 | 0.3 |
|  | 443 | 36 | -71 | -16 | 10.8 | 0.3 |
|  | 447 | 29 | -47 | -14 | 8.3 | 0.3 |
|  | 450 | 41 | -48 | -20 | 8.4 | 0.3 |
|  | 452 | 28 | -59 | -12 | 9.7 | 0.4 |
| Postcentral | 476 | 60 | -18 | 37 | 9.2 | 0.3 |
|  | 478 | 42 | -31 | 49 | 8.5 | 0.3 |
|  | 484 | 30 | -40 | 61 | 8.5 | 0.2 |
| Parietal (superior) | 494 | -26 | -61 | 61 | 9.1 | 0.1 |
|  | 495 | -27 | -53 | 68 | 8 | 0.1 |
|  | 497 | -21 | -67 | 47 | 9.3 | 0.2 |
|  | 498 | -15 | -69 | 50 | 8.5 | 0.1 |
|  | 499 | -28 | -69 | 50 | 8.9 | 0.3 |
|  | 501 | -17 | -79 | 50 | 8.7 | 0.1 |
|  | 502 | 30 | -60 | 64 | 8.8 | 0.2 |
|  | 504 | 34 | -62 | 57 | 9.3 | 0.3 |
|  | 506 | 33 | -50 | 58 | 9.8 | 0.3 |
|  | 507 | 21 | -57 | 74 | 8.1 | 0.1 |
|  | 509 | 31 | -73 | 53 | 8.6 | 0.1 |
|  | 510 | 21 | -74 | 55 | 8.8 | 0.1 |
|  | 511 | 20 | -65 | 54 | 9.2 | 0.2 |
| Parietal (inferior) | 514 | -45 | -30 | 42 | 8.6 | 0.2 |
|  | 516 | -32 | -75 | 44 | 8 | 0.2 |
|  | 521 | -39 | -47 | 42 | 8.6 | 0.2 |
|  | 522 | -32 | -46 | 50 | 8.5 | 0.1 |
|  | 523 | -31 | -53 | 46 | 8.4 | 0.2 |
|  | 527 | 46 | -39 | 50 | 8.6 | 0.4 |
|  | 529 | 37 | -49 | 46 | 8.7 | 0.3 |
|  | 530 | 35 | -44 | 51 | 9.2 | 0.3 |
| Supramarginal | 536 | -61 | -28 | 34 | 8.4 | 0.2 |
|  | 539 | 44 | -34 | 41 | 8.7 | 0.2 |
|  | 542 | 63 | -24 | 37 | 8.5 | 0.2 |
| Angular | 557 | 34 | -60 | 44 | 8.7 | 0.4 |
| Precuneus | 561 | -5 | -77 | 53 | 7.9 | 0.1 |
|  | 573 | -9 | -71 | 54 | 8.1 | 0.1 |
|  | 576 | 14 | -71 | 45 | 7.9 | 0.1 |
| Temporal (middle) | 678 | -45 | -67 | 11 | 8.9 | 0.1 |
|  | 685 | -49 | -62 | 0 | 9.3 | 0.2 |
|  | 701 | 52 | -59 | 3 | 8.9 | 0.2 |
|  | 717 | 49 | -69 | 4 | 10.5 | 0.2 |
| Temporal (inferior) | 728 | -54 | -58 | -11 | 8.5 | 0.3 |
|  | 732 | -45 | -52 | -13 | 9.3 | 0.3 |
|  | 755 | 46 | -53 | -11 | 9.7 | 0.4 |

## Supplement

**Figure S1:**
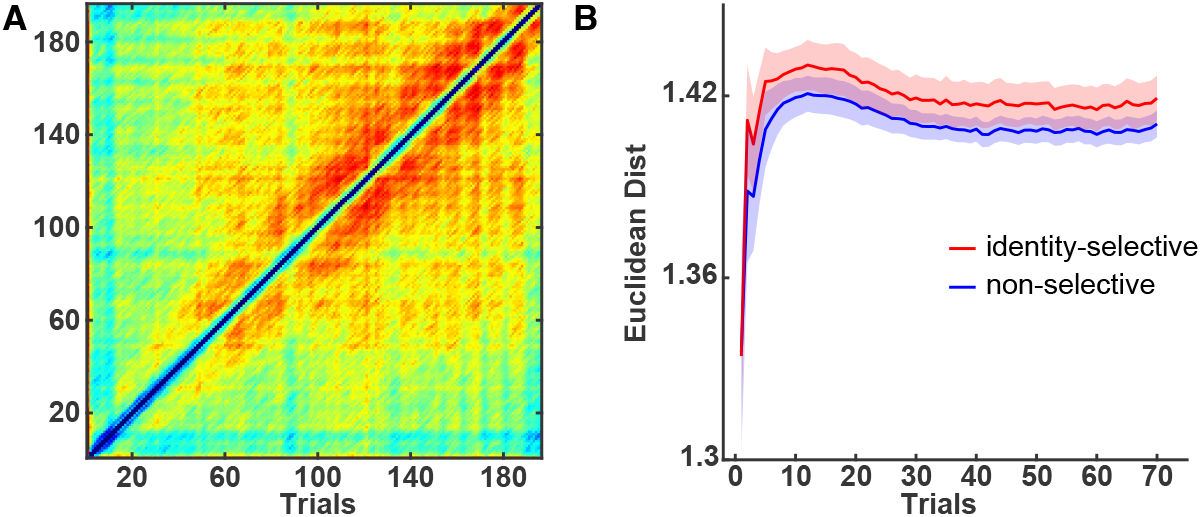
Pairwise distance between responses as a function of delay. Pairwise response distances *d*_*w,u,r*_ (*i, j*) between trials *i* and *j* in parcel *w*, run *r* and subject *u* were averaged over subjects and runs to obtain *T*_*w*_ (*i, j*) = *(d*_*w,u,r*_ (*i, j*))_*u,r*_ **A)** Pattern of response distances for all pairs of trials *i, j*, average *(T*_*w*_ (*i, j*) ) _*w*_ over all parcels. **B**) Pattern of response distances as a function of delay, |*i −j*|, average (±SEM) over identity-selective parcels (red) and non-identity-selective parcels (blue).

**Figure S2:**
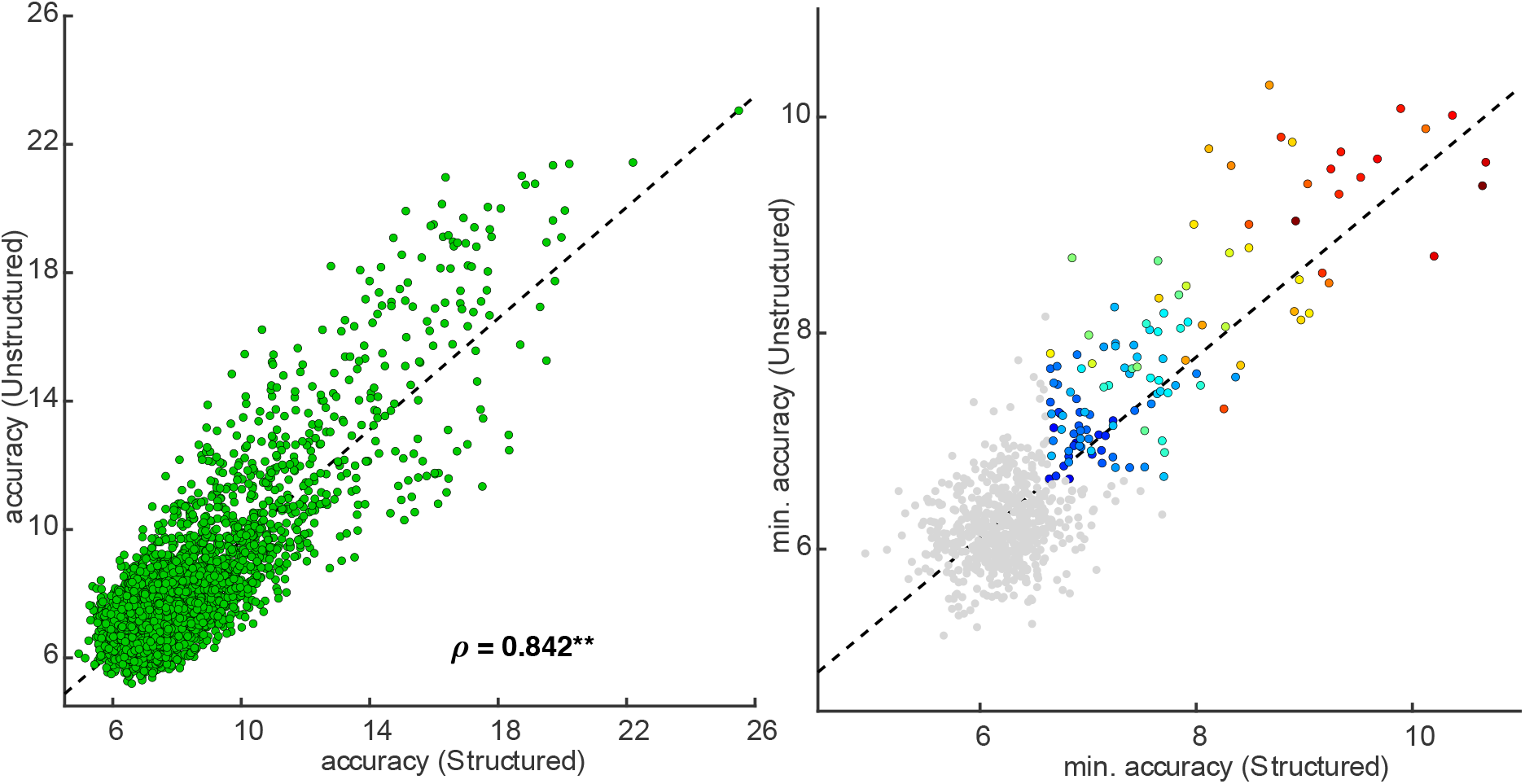
Classification accuracy *a*_*w,u*_ for all observers *u* and parcels *w*, as observed with structured or with unstructured presentation sequences. **A)** Accuracy values observed with structured and unstructured sequences are correlated (*ρ* = 0.842, *p* < 0.001). **B)** Minimal accuracy values of all parcels *w*, obtained by combining observers. Values from structured and unstructured sequences are correlated (*ρ* = 0.802, *p* < 0.001). Colors indicate the combined accuracy of parcels categorized as identity-selective.

**Figure S3:**
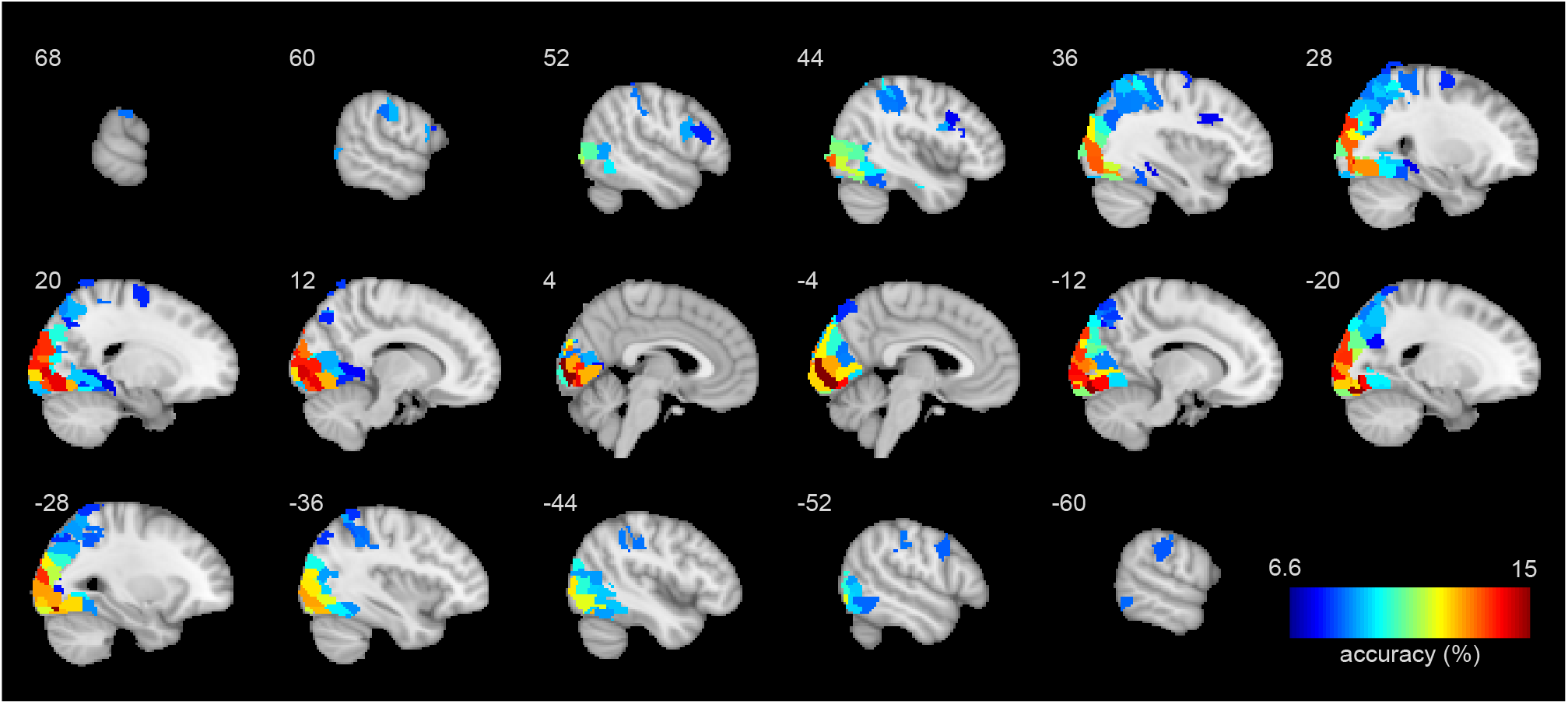
Cortical representation of object ‘identity’. Color scale indicates average classification accuracy of identity-selective parcels, ranging from chance level (6.67%) to maximum (15%). Sagittal slices (8 mm thickness) range from *X* = *−*60 to *X* = +68 (MNI).

**Figure S4:**
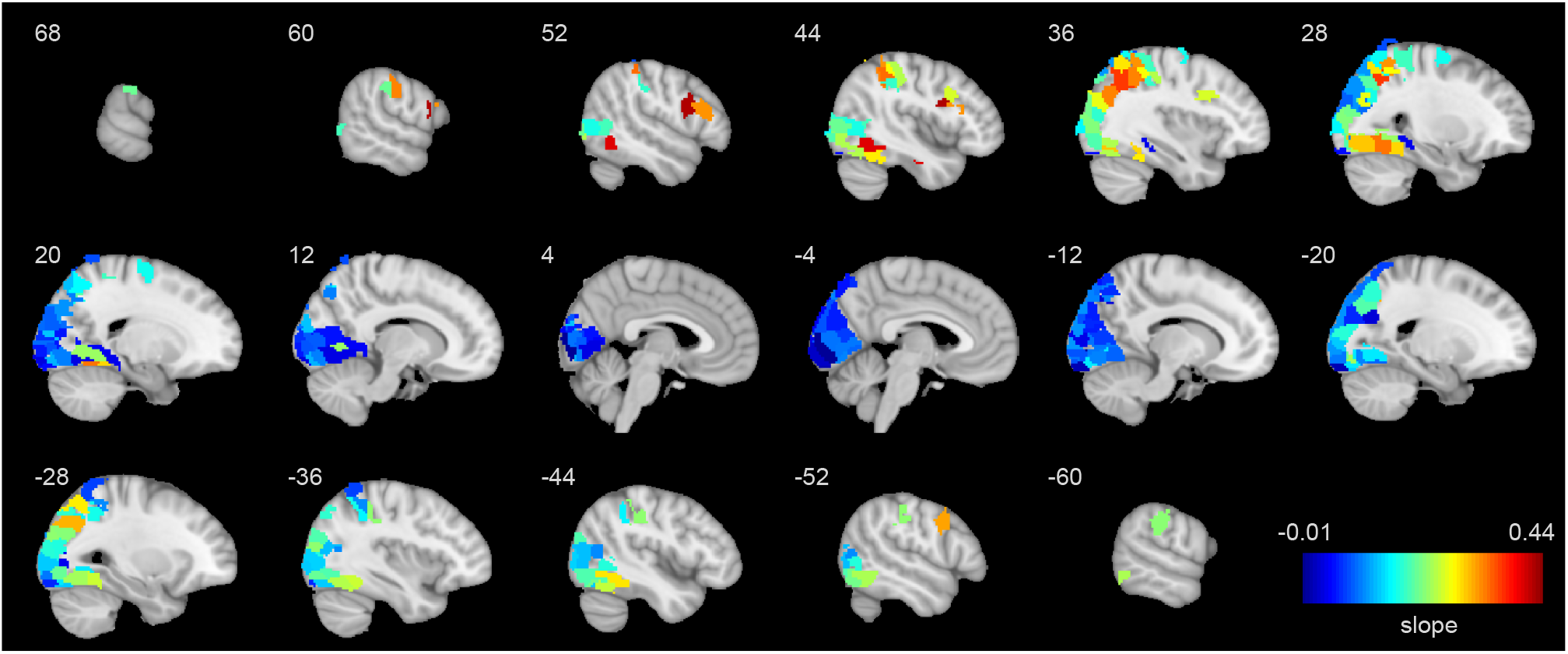
Cortical representation of object ‘novelty’. Color scale indicates the average rate *β*^*novelty*^ of increase in the variance ratio *F*^*novelty*^, ranging from a minimal value of *−*0.01 to a maximal value of 0.44. Sagittal slices (8 mm thickness) range from *X* = *−*60 to *X* = +68 (MNI).

**Figure S5:**
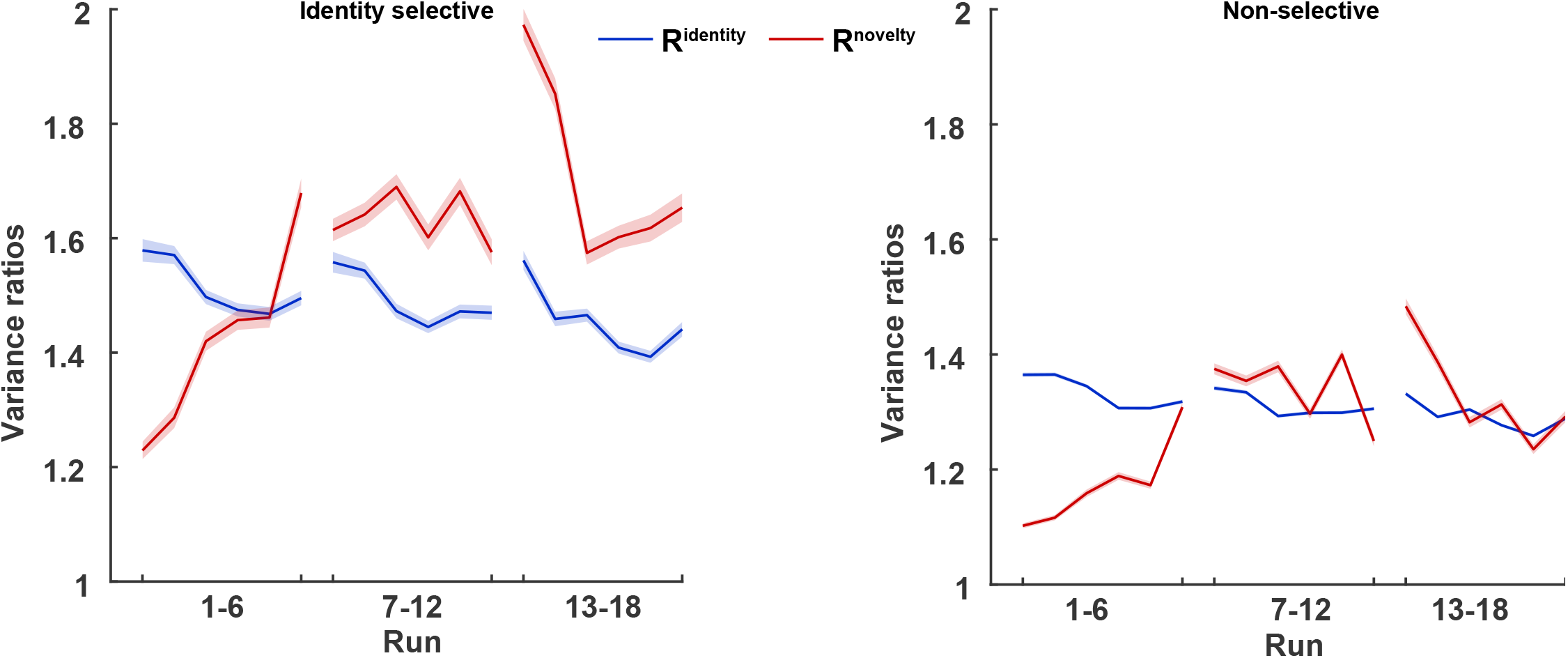
Development of variance fractions (mean ± *S*.*E*.*M*) over the course of experiment (18 runs in 3 session). Variance ratios *F*^*novelty*^ (red) and *F*^*identity*^ (blue) were computed for every run, and averaged separately for identity-selective (left) and non-identity-selective parcels (right).

## References

C. Allefeld, K. GÖrgen, and J.-D. Haynes. Valid population inference for information-based imaging: From the second-level t-test to prevalence inference. Neuroimage, 141:378–392, 2016. doi:10.1016/j.neuroimage.2016.07.040.

M. J. Anderson. A new method for non-parametric multivariate analysis of variance. Austral Ecology, 26(1):32–46, 2001. doi:10.1111/j.1442-9993.2001.01070.pp.x.

C. F. Beckmann and S. M. Smith. Probabilistic independent component analysis for functional magnetic resonance imaging. IEEE transactions on medical imaging, 23(2):137–152, 2004. doi:10.1109/TMI.2003.822821.

Y. Bi, X. Wang, and A. Caramazza. Object domain and modality in the ventral visual pathway. Trends in Cognitive Sciences, 20(4):282–290, 2016. doi:10.1016/j.tics.2016.02.002.

D. H. Brainard. The psychophysics toolbox. Spatial Vision, 10(4):433–436, 1997. doi:10.1163/156856897X00357.

M. Brants, J. Wagemans, and H. P. Op de Beeck. Activation of fusiform face area by greebles is related to face similarity but not expertise. Journal of Cognitive Neuroscience, 23(12): 3949–3958, 2011. doi:10.1162/jocna00072.

M. Brants, J. Bulthé, N. Daniels, J. Wagemans, and H. P. O. de Beeck. How learning might strengthen existing visual object representations in human object-selective cortex. Neuroimage, 127:74–85, 2016. doi:10.1016/j.neuroimage.2015.11.063.

C. M. Bukach, I. Gauthier, and M. J. Tarr. Beyond faces and modularity: the power of an expertise framework. Trends in Cognitive Sciences, 10(4):159–166, 2006. doi:10.1016/j.tics.2006.02.004.

J. S. Cetron, A. C. Connolly, S. G. Diamond, V. V. May, J. V. Haxby, and D. J. Kraemer. Decoding individual differences in stem learning from functional mri data. Nature Communications, 10(1):1–10, 2019. doi:10.1038/s41467-019-10053-y.

I. Charest and N. Kriegeskorte. The brain of the beholder: honouring individual representational idiosyncrasies. Language, Cognition and Neuroscience, 30(4):367–379, 2015. doi:10.1080/23273798.2014.1002505.

T. B. Christophel, P. C. Klink, B. Spitzer, P. R. Roelfsema, and J.-D. Haynes. The distributed nature of working memory. Trends in Cognitive Sciences, 21(2):111–124, 2017. doi:10.1016/j.tics.2016.12.007.

E. Collins and M. Behrmann. Exemplar learning reveals the representational origins of expert category perception. Proceedings of the National Academy of Sciences, USA, 117(20):11167– 11177, 2020. doi:10.1073/pnas.1912734117.

A. C. Connolly, J. S. Guntupalli, J. Gors, M. Hanke, Y. O. Halchenko, Y.-C. Wu, H. Abdi, and J. V. Haxby. The representation of biological classes in the human brain. Journal of Neuroscience, 32(8):2608–2618, 2012. doi:10.1523/JNEUROSCI.5547-11.2012.

H. P. O. de Beeck and C. I. Baker. The neural basis of visual object learning. Trends in Cognitive Sciences, 14(1):22–30, 2010. doi:10.1016/j.tics.2009.11.002.

H. P. O. de Beeck, C. I. Baker, J. J. DiCarlo, and N. G. Kanwisher. Discrimination training alters object representations in human extrastriate cortex. Journal of Neuroscience, 26(50): 13025–13036, 2006. doi:10.1523/JNEUROSCI.2481-06.2006.

J. V. Dornas and J. Braun. Finer parcellation reveals detailed correlational structure of resting-state fmri signals. Journal of Neuroscience Methods, 294:15–33, 2018. doi:10.1016/j.jneumeth.2017.10.020.

S. Duyck, F. Martens, C.-Y. Chen, and H. Op de Beeck. How visual expertise changes representational geometry: A behavioral and neural perspective. Journal of Cognitive Neuroscience, 33(12):2461–2476, 2021. doi:10.1162/jocna01778.

E. Eger, J. Ashburner, J.-D. Haynes, R. J. Dolan, and G. Rees. fmri activity patterns in human loc carry information about object exemplars within category. Journal of Cognitive Neuroscience, 20(2):356–370, 2008. doi:10.1162/jocn.2008.20019.

E. Freud, J. C. Culham, D. C. Plaut, and M. Behrmann. The large-scale organization of shape processing in the ventral and dorsal pathways. eLife, 6:e27576, 2017. doi:10.7554/eLife.34464.

I. Gauthier and M. J. Tarr. Visual object recognition: Do we (finally) know more now than we did? Annual Review of Vision Science, 2:377–396, 2016. doi:10.1146/annurev-vision-111815-114621.

I. Gauthier, M. J. Tarr, A. W. Anderson, P. Skudlarski, and J. C. Gore. Activation of the middle fusiform’face area’increases with expertise in recognizing novel objects. Nature Neuroscience, 2(6):568–573, 1999. doi:10.1038/9224.

D. N. Greve and B. Fischl. Accurate and robust brain image alignment using boundary-based registration. Neuroimage, 48(1):63–72, 2009. doi:10.1016/j.neuroimage.2009.06.060.

K. Grill-Spector and K. S. Weiner. The functional architecture of the ventral temporal cortex and its role in categorization. Nature Reviews Neuroscience, 15(8):536–548, 2014. doi:10.1038/nrn3747.

K. Grill-Spector, N. Knouf, and N. Kanwisher. The fusiform face area subserves face perception, not generic within-category identification. Nature Neuroscience, 7(5):555–562, 2004. doi:10.1038/nn1224.

A. Harel, D. Kravitz, and C. I. Baker. Beyond perceptual expertise: revisiting the neural substrates of expert object recognition. Frontiers in Human Neuroscience, 7:885, 2013. doi:10.1167/14.10.820.

J. V. Haxby. Multivariate pattern analysis of fmri: the early beginnings. Neuroimage, 62(2): 852–855, 2012. doi:10.1016/j.neuroimage.2012.03.016.

J. V. Haxby, M. I. Gobbini, M. L. Furey, A. Ishai, J. L. Schouten, and P. Pietrini. Distributed and overlapping representations of faces and objects in ventral temporal cortex. Science, 293 (5539):2425–2430, 2001. doi:10.1126/science.1063736.

C. P. Hung, G. Kreiman, T. Poggio, and J. J. DiCarlo. Fast readout of object identity from macaque inferior temporal cortex. Science, 310(5749):863–866, 2005. doi:10.1126/science.1117593.

M. Jenkinson and S. Smith. A global optimisation method for robust affine registration of brain images. Medical Image Analysis, 5(2):143–156, 2001. doi:10.1016/S1361-8415(01)00036-6.

M. Jenkinson, P. Bannister, M. Brady, and S. Smith. Improved optimization for the robust and accurate linear registration and motion correction of brain images. Neuroimage, 17(2): 825–841, 2002. doi:10.1006/nimg.2002.1132.

S. K. Jeong and Y. Xu. Behaviorally relevant abstract object identity representation in the human parietal cortex. Journal of Neuroscience, 36(5):1607–1619, 2016. doi:10.1523/JNEUROSCI.1016-15.2016.

E. Kakaei, S. Aleshin, and J. Braun. Visual object recognition is facilitated by temporal community structure. Learning & Memory, 28(5):148–152, 2021. doi:10.1101/lm.053306.120.

C. S. Konen and S. Kastner. Two hierarchically organized neural systems for object information in human visual cortex. Nature Neuroscience, 11(2):224–231, 2008. doi:10.1038/nn2036.

T. Konkle and A. Oliva. A real-world size organization of object responses in occipitotemporal cortex. Neuron, 74(6):1114–1124, 2012. doi:10.1016/j.neuron.2012.04.036.

D. J. Kravitz, K. S. Saleem, C. I. Baker, and M. Mishkin. A new neural framework for visuospatial processing. Nature Reviews Neuroscience, 12(4):217–230, 2011. doi:10.1038/nrn3008.

D. J. Kravitz, K. S. Saleem, C. I. Baker, L. G. Ungerleider, and M. Mishkin. The ventral visual pathway: an expanded neural framework for the processing of object quality. Trends in Cognitive Sciences, 17(1):26–49, 2013. doi:10.1016/j.tics.2012.10.011.

N. Kriegeskorte and J. Diedrichsen. Peeling the onion of brain representations. Annual Review of Neuroscience, 42:407–432, 2019. doi:10.1146/annurev-neuro-080317-061906.

N. Kriegeskorte, R. Goebel, and P. Bandettini. Information-based functional brain mapping. Proceedings of the National Academy of Sciences, USA, 103(10):3863–3868, 2006. doi:10.1073/pnas.0600244103.

N. Kriegeskorte, M. Mur, and P. A. Bandettini. Representational similarity analysis-connecting the branches of systems neuroscience. Frontiers in Systems Neuroscience, 2:4, 2008a. doi:10.3389/neuro.06.004.2008.

N. Kriegeskorte, M. Mur, D. A. Ruff, R. Kiani, J. Bodurka, H. Esteky, K. Tanaka, and P. A. Bandettini. Matching categorical object representations in inferior temporal cortex of man and monkey. Neuron, 60(6):1126–1141, 2008b. doi:10.1016/j.neuron.2008.10.043.

H. Liu, Y. Agam, J. R. Madsen, and G. Kreiman. Timing, timing, timing: fast decoding of object information from intracranial field potentials in human visual cortex. Neuron, 62(2): 281–290, 2009. doi:10.1016/j.neuron.2009.02.025.

F. Martens, J. Bulthé, C. van Vliet, and H. O. de Beeck. Domain-general and domain-specific neural changes underlying visual expertise. Neuroimage, 169:80–93, 2018. doi:10.1016/j.neuroimage.2017.12.013.

R. W. McGugin, J. C. Gatenby, J. C. Gore, and I. Gauthier. High-resolution imaging of expertise reveals reliable object selectivity in the fusiform face area related to perceptual performance. Proceedings of the National Academy of Sciences, USA, 109(42):17063–17068, 2012. doi:10.1073/pnas.1116333109.

A. Nestor, D. C. Plaut, and M. Behrmann. Feature-based face representations and image reconstruction from behavioral and neural data. Proceedings of the National Academy of Sciences, USA, 113(2):416–421, 2016. doi:10.1073/pnas.1514551112.

A. X. Patel, P. Kundu, M. Rubinov, P. S. Jones, P. E. Vértes, K. D. Ersche, J. Suckling, and E. T. Bullmore. A wavelet method for modeling and despiking motion artifacts from restingstate fmri time series. Neuroimage, 95:287–304, 2014. doi:10.1016/j.neuroimage.2014.03.012.

C. J. Perry and M. Fallah. Feature integration and object representations along the dorsal stream visual hierarchy. Frontiers in Computational Neuroscience, 8:84, 2014. doi:10.3389/fncom.2014.00084.

C. C. Poirier, A. G. De Volder, D. Tranduy, and C. Scheiber. Neural changes in the ventral and dorsal visual streams during pattern recognition learning. Neurobiology of Learning and Memory, 85(1):36–43, 2006. doi:10.1016/j.nlm.2005.08.006.

R. A. Poldrack. Imaging brain plasticity: conceptual and methodological issuesa theoretical review. Neuroimage, 12(1):1–13, 2000. doi:10.1006/nimg.2000.0596.

Z. N. Roth and E. Zohary. Fingerprints of learned object recognition seen in the fmri activation patterns of lateral occipital complex. Cerebral Cortex, 25(9):2427–2439, 2015. doi:10.1093/cercor/bhu042.

S. M. Smith. Fast robust automated brain extraction. Human Brain Mapping, 17(3):143–155, 2002. doi:10.1002/hbm.10062.

S. M. Smith and J. M. Brady. Susana new approach to low level image processing. International Journal of Computer Vision, 23(1):45–78, 1997. doi:10.1023/A:1007963824710.

N. Tzourio-Mazoyer, B. Landeau, D. Papathanassiou, F. Crivello, O. Etard, N. Delcroix, B. Mazoyer, and M. Joliot. Automated anatomical labeling of activations in spm using a macroscopic anatomical parcellation of the mni mri single-subject brain. Neuroimage, 15 (1):273–289, 2002. doi:10.1006/nimg.2001.0978.

L. Q. Uddin. Salience processing and insular cortical function and dysfunction. Nature Reviews Neuroscience, 16(1):55–61, 2015. doi:10.1038/nrn3857.

M. Visconti di Oleggio Castello, J. V. Haxby, and M. I. Gobbini. Shared neural codes for visual and semantic information about familiar faces in a common representational space. Proceedings of the National Academy of Sciences, USA, 118(45):e2110474118, 2021. doi:10.1073/pnas.2110474118.

L. Wang, R. E. Mruczek, M. J. Arcaro, and S. Kastner. Probabilistic maps of visual topography in human cortex. Cerebral Cortex, 25(10):3911–3931, 2015. doi:10.1093/cercor/bhu277.

K. S. Weiner and K. Zilles. The anatomical and functional specialization of the fusiform gyrus. Neuropsychologia, 83:48–62, 2016. doi:10.1016/j.neuropsychologia.2015.06.033.

A. C.-N. Wong, T. J. Palmeri, and I. Gauthier. Conditions for facelike expertise with objects: Becoming a ziggerin expertbut which type? Psychological Science, 20(9):1108–1117, 2009. doi:10.1111/j.1467-9280.2009.02430.x.

Y. K. Wong, J. R. Folstein, and I. Gauthier. The nature of experience determines object representations in the visual system. Journal of Experimental Psychology: General, 141(4): 682, 2012. doi:10.1037/a0027822.

M. F. Wurm and A. Caramazza. Two whatpathways for action and object recognition. Trends in Cognitive Sciences, 2021. doi:10.1016/j.tics.2021.10.003.

J. Ye, T. Xiong, and D. Madigan. Computational and theoretical analysis of null space and orthogonal linear discriminant analysis. Journal of Machine Learning Research, 7(7), 2006. URL http://jmlr.org/papers/v7/ye06a.html.

I. Yildirim, J. Wu, N. Kanwisher, and J. Tenenbaum. An integrative computational architecture for object-driven cortex. Current Opinion in Neurobiology, 55:73–81, 2019. doi:10.1016/j.conb.2019.01.010.

H. Yu and J. Yang. A direct lda algorithm for high-dimensional datawith application to face recognition. Pattern Recognition, 34(10):2067–2070, 2001. doi:10.1016/S0031-3203(00)00162-X.

X. Yue, B. S. Tjan, and I. Biederman. What makes faces special? Vision Research, 46(22): 3802–3811, 2006. doi:10.1016/j.visres.2006.06.017.

T. P. Zanto, M. T. Rubens, J. Bollinger, and A. Gazzaley. Top-down modulation of visual feature processing: the role of the inferior frontal junction. Neuroimage, 53(2):736–745, 2010. doi:10.1016/j.neuroimage.2010.06.012.

T. P. Zanto, M. T. Rubens, A. Thangavel, and A. Gazzaley. Causal role of the prefrontal cortex in top-down modulation of visual processing and working memory. Nature Neuroscience, 14 (5):656–661, 2011. doi:10.1038/nn.2773.

Y. Zhang, M. Brady, and S. Smith. Segmentation of brain mr images through a hidden markov random field model and the expectation-maximization algorithm. IEEE Transactions on Medical Imaging, 20(1):45–57, 2001. doi:10.1109/42.906424.

